# Multi-omics data of a weedy coral species from Ulithi Atoll (Micronesia) to investigate the impact of human disturbance on coral health and resilience

**DOI:** 10.64898/2026.05.21.726996

**Authors:** Erin E. Chille, Gabriella M. Panayotakis, Timothy G. Stephens, Michelle Paddack, Nicole L. Crane, Giacomo Bernardi, John Rulmal, Debashish Bhattacharya

**Affiliations:** Department of Biochemistry and Microbiology, Rutgers University, New Brunswick, New Jersey, USA; One People One Reef, Santa Cruz, California, USA; Santa Barbara City College, Santa Barbara, California, USA; David H. Smith Conservation Research Fellowship, Society for Conservation Biology, Washington, DC, USA; Department of Ecology and Evolutionary Biology, University of California, Santa Cruz, California, USA; Ulithi Falalop Community Action Program, Yap, FSM

**Keywords:** adaptation, anthropogenic stress, biomarkers, chronic disturbance, corals, genomics, metabolomics, multi-omics, proteomics, urbanization

## Abstract

**Background:** Coral reefs are among the most biodiverse ecosystems on Earth yet face an increasing risk of collapse under climate change and intensifying anthropogenic pressures. Although there is strong evidence that stressors such as overfishing, sedimentation, and wastewater runoff can drive shifts in coral community structure, few studies have investigated the molecular mechanisms underlying these shifts. Integrating data across multiple layers of biological organization, from the genome to the proteome and metabolome, offers a powerful approach to understanding how coral physiology and resilience are impacted by human activities.

**Findings:** Here, we present high-coverage whole genome sequencing (*Montipora* only) together with untargeted metabolomic and proteomic datasets from three coral genera (*Acropora*, *Pocillopora*, and *Montipora*) that inhabit Ulithi Atoll, Yap State, FSM. Ulithi Atoll is an ideal system for investigating how anthropogenic pressures influence coral molecular ecology and adaptation, with extensive historical ecological data available through the long-term research program of the One People One Reef collaborative to connect multi-omics data with macroecological trends. Following data cleaning and quality control, all datasets exhibited minimal technical variation among samples, providing a comprehensive baseline for future research on coral resilience across a mosaic of natural and anthropogenic stressors.

**Conclusions:** These data provide a resource for examining how anthropogenic pressures influence coral physiology and resilience across environmental gradients. They also allow the identification of biomarkers of chronic stress that may inform the development of point-of-care diagnostic tools for coral health and contribute to evidence-based reef conservation and management strategies.

## Context

Under global climate change, the resilience of an ecosystem and its capacity to absorb and then recover from disturbance, is becoming increasingly recognized as a critical management priority [1]. In coral reef ecosystems, resilience largely depends on the health and resilience of coral populations [2]. As foundation species, reef-building corals create complex habitats that drive biological diversity and provide life-sustaining ecosystem services to humans, such as fisheries and coastal protection [3]. Understanding how environmental change and anthropogenic activities affect coral health at the population level is therefore a priority for evidence-based coral reef conservation and management.

Coral health depends on the interplay between genetic variation, environmental factors, and the host-associated microbes that make up the coral holobiont, of which core members are single-celled algal endosymbionts in the dinoflagellate family, Symbiodiniaceae. This mutualistic symbiosis is obligate for the host and “powers” tropical coral reef ecosystems through the exchange of sugars, lipids, and other metabolites between the cnidarian animal and algal endosymbionts [4]. To date, coral physiology and health have been primarily studied through the lens of thermal stress and bleaching (loss of pigments in and/or expulsion of the Symbiodiniaceae). However, local stressors from anthropogenic activity, including sedimentation, sewage discharge, eutrophication, and overfishing of herbivorous fishes, may also exacerbate the risk of coral mortality [5–7] and alter reef community composition [8–11].

Although these pressures have clear impacts on reef community assembly [8–11], their effects on the coral holobiont at the molecular level remain poorly characterized. Integrating data across multiple layers of biological organization, including the genome, proteome, and metabolome offers more holistic insights into the impacts of chronic human disturbance on coral holobiont molecular ecology and adaptation. These shifts are not easily discerned visually and allow the identification of biomarkers that can be incorporated into point-of-care tools for ecosystem management [12].

Ulithi Atoll, a 500 km^2^ reef system in Yap State, Federated States of Micronesia, is an ideal system to investigate how environmental and anthropogenic factors influence coral biology. Whereas human population size has historically been low (< 1,000 people), and communities maintain a largely traditional way of life, it has nonetheless influenced the reef ecosystem through resource use, infrastructure development, modern fishing technologies, and high military activity during World War II [8]. Human settlements are largely situated on the northern and eastern side of the atoll, generating spatial gradients in anthropogenic activities. Importantly, these environmental differences are decoupled from genetic connectivity, which is strongest East-West [13], providing the opportunity to examine how environmental conditions shape coral molecular phenotypes independent of strong population genetic structure.

Ecological surveys conducted at Ulithi demonstrate that reef community composition is structured along gradients of human presence and environmental exposure [8]. Hierarchical clustering of benthic and fish community composition identified three major ecological clusters (**Figure 1**): oceanic reefs near uninhabited islands (blue), oceanic reefs near inhabited islands (purple), and lagoon-facing reefs (red; **Figure 1**). Clusters are predicted by village proximity, population size, and reef exposure to deeper water [8]. Sites with the highest levels of human impact have experienced a shift in the coral community assemblage, with a reef-building coral in the genus *Montipora* (hereinafter referred to as *Montipora* sp. 1. *aff. capitata*) rapidly colonizing disturbed (e.g., due to anthropogenic activities, typhoons) lagoonal reef zones [8,14]. This species is outcompeting many functionally important reef-building corals, such as *Pocillopora* and *Acropora,* which host a rich diversity of food fish, raising the question of how different coral taxa respond at the molecular level to chronic human disturbance.

**Figure 1.**
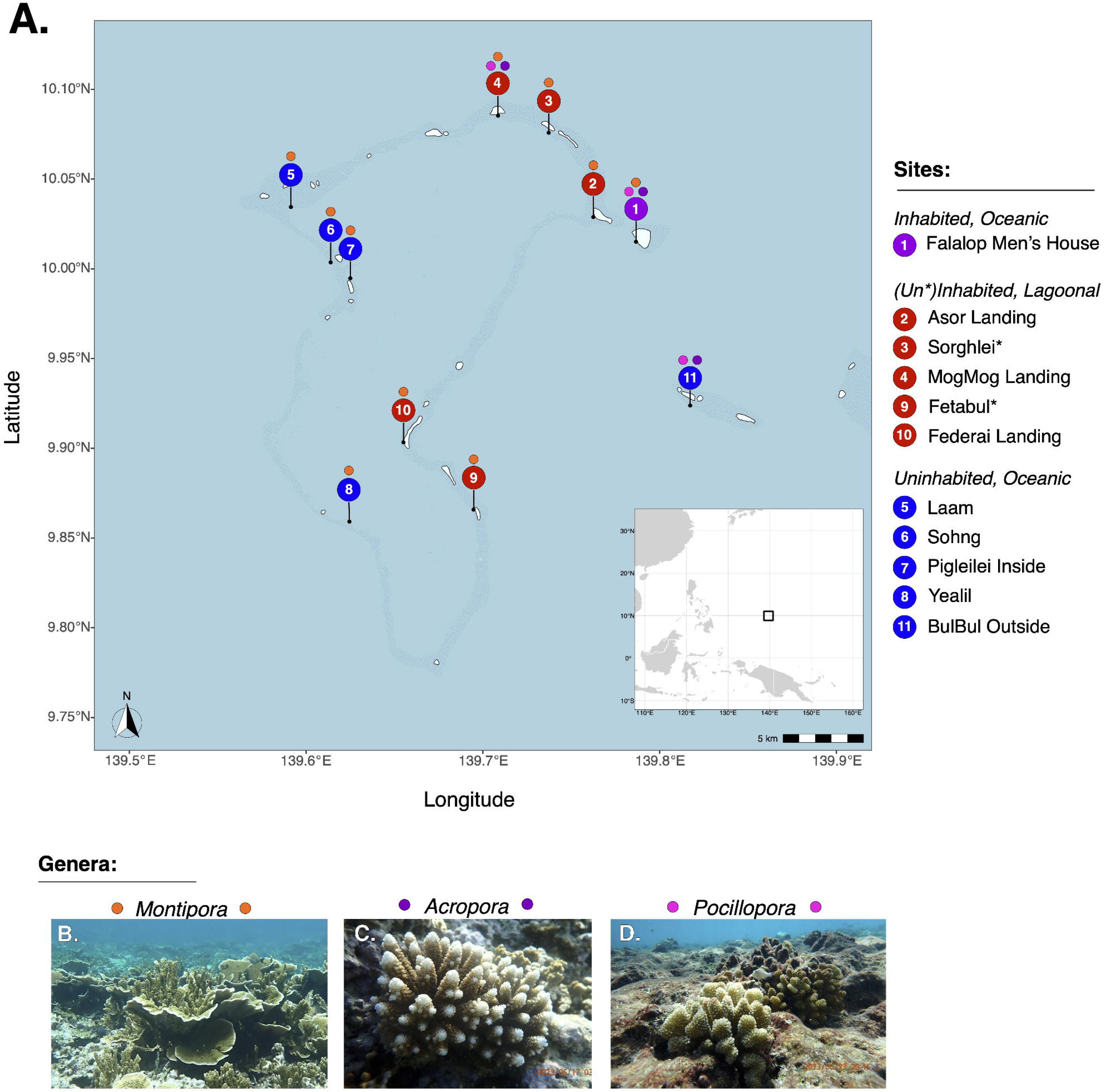
Sampling design and study sites at Ulithi Atoll. **(A)** Map of sampling locations across Ulithi Atoll (Yap State, Federated States of Micronesia). Site colors correspond to the ecological clusters identified by Crane et al. [8] based on benthic community composition. See **Table 1** for site metadata and coordinates. Colored circles above each site marker indicate the coral genera sampled at that location (*Montipora*, orange; *Acropora*, purple; *Pocillopora*, magenta; representative images for each genus shown at the bottom of the figure). At each site, 10 colonies per genus were collected, except where noted in **Table S1**. **(B-D)** Representative colonies of the sampled genera: **(B)** *Montipora*, **(C)** *Acropora*, and **(D)** *Pocillopora*. Coral images taken by EEC. *Indicates an uninhabited lagoonal site that ecologically clustered with the inhabited, lagoonal sites in Crane et al. [9].

**Table 1.**
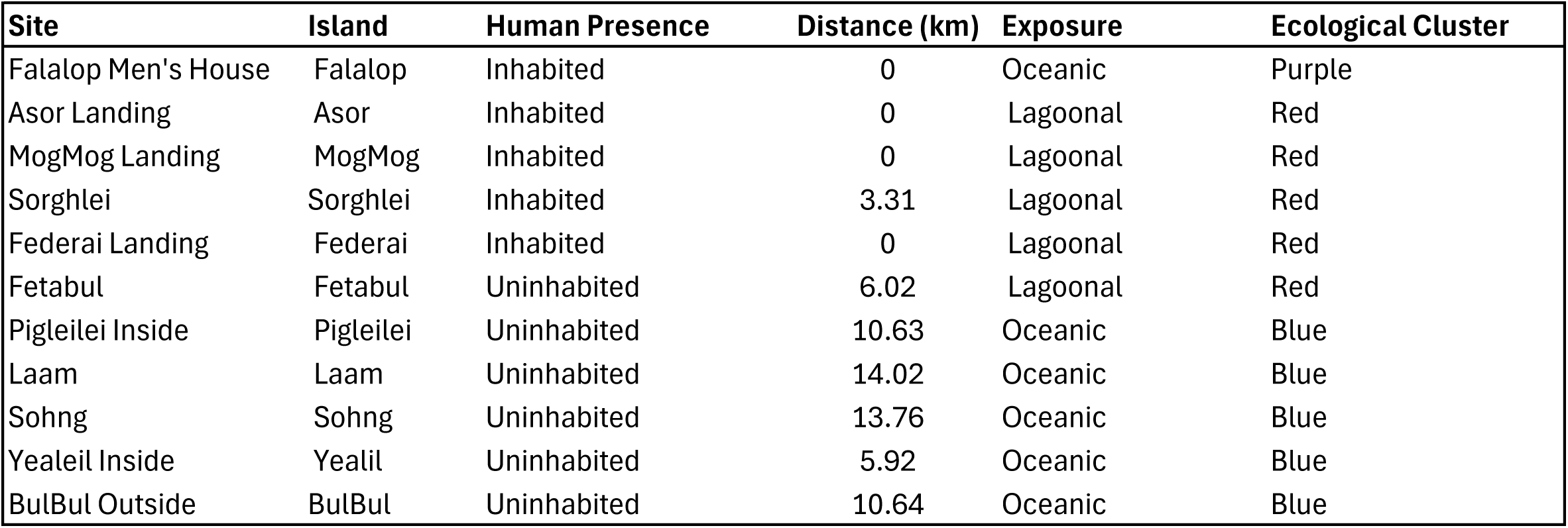
Summary of collection site metadata. Columns from left to right correspond to: site name, the island where the site s located, the island(s) with jurisdiction for reef and fishery management, human presence (inhabited/uninhabited during the study period), straight line distance from the closest jurisdiction, whether the site was located within the Ulithi lagoon or exposed directly to the ocean, and ecological cluster defined from hierarchical clustering of benthic community data from Crane et al. [8] as it appears in **Fig. 1**.

| Site | Island | Human Presence | Distance (km) | Exposure | Ecological Cluster |
| --- | --- | --- | --- | --- | --- |
| Falalop Men's House | Falalop | Inhabited | 0 | Oceanic | Purple |
| Asor Landing | Asor | Inhabited | 0 | Lagoonal | Red |
| MogMog Landing | MogMog | Inhabited | 0 | Lagoonal | Red |
| Sorghlei | Sorghlei | Inhabited | 3.31 | Lagoonal | Red |
| Federali Landing | Federali | Inhabited | 0 | Lagoonal | Red |
| Fetabul | Fetabul | Uninhabited | 6.02 | Lagoonal | Red |
| Pigleilei Inside | Pigleilei | Uninhabited | 10.63 | Oceanic | Blue |
| Laam | Laam | Uninhabited | 14.02 | Oceanic | Blue |
| Sohng | Sohng | Uninhabited | 13.76 | Oceanic | Blue |
| Yealeil Inside | Yealil | Uninhabited | 5.92 | Oceanic | Blue |
| BulBul Outside | BulBul | Uninhabited | 10.64 | Oceanic | Blue |

Here, we describe the generation of genomic (*Montipora* only), proteomic, and metabolomic datasets generated from coral colonies sampled at different sites in Ulithi Atoll. By integrating these datasets with ecological survey data [8], photographic documentation of sampled colonies (when available), and associated environmental data (**Table 1**), this resource supports the investigation of how anthropogenic activities influence coral molecular ecology and adaptation across multiple omics layers, including genetic variation, protein expression, and metabolomic profiles. Samples include representatives from *Montipora* sp. 1. *aff. capitata*, *Pocillopora* spp., and *Acropora* spp., which will enable comparative analyses between taxa.

## Methods

### Sample collection

#### Ethics statement

We received specific permission to sample reefs from Philip Paiy, Chief of Falalop, Lipipi Clan, representing Council of Ten (CO-X), Ulithi Atoll, and from the Council of Tamol – representing the outer islands of Yap and the Yap legislature. We also received support from Yap Marine Resources. We had formal meetings with the Governor’s office (Yap) and the Legislature to brief them on the work. Finally, we received on-site permission from village Chiefs and/or reef owners (or designee) prior to each sampling visit before any in-water work was conducted.

#### Sampling methods

*Montipora* (*n* = 100)*, Pocillopora* (*n* = 25), and *Acropora* (*n* = 26) samples were collected from 10 long-term sampling sites in Ulithi Atoll, Yap State, Federated States of Micronesia (*n* ≥ 5 colonies/site; **Table 2**). To ensure even sampling across the sampling sites, two 50 meter transects were laid out at each site and samples were collected from colonies lying directly beneath the transect line every 10 meters. At sites where the density of colonies was too low for this approach (e.g., Falalop Men’s House, BulBul Outside), colonies were opportunistically sampled and tagged until collection was complete to prevent accidental re-sampling. When possible, each coral colony was photographed immediately prior to sampling. We sampled nubbins (∼1 g) from each coral colony. After both transects were sampled, nubbins were split into 2 mL tubes for DNA, metabolite, and protein preservation. For DNA and metabolite preservation, each sample was placed in a tube containing 0.5 mm glass beads (Millipore Sigma cat #19-622) and either 1 mL DNA/RNA Shield for nucleic acid preservation (Zymo Research cat #R1100) or 1 mL OMNImet·GUT metabolite stabilizing solution (DNA Genotek Catalog #ME-200). The nubbins in these preservation tubes were lysed for 5 minutes using a vortex mixer set to maximum speed. For protein preservation, samples were placed in 1 mL of AllProtect Tissue Reagent (Qiagen cat #76405). Samples were stored in the preservative in a cooler with ice packs during collection, stored at −20°C in the field, shipped on ice packs, and transferred to −80° in the laboratory until sample processing.

**Table 2.** Summary of *Montipora sp. 1 aff. capitata* samples collected. Columns from left to right correspond to: genus, sample name, sample id, collection date, site name, the position on the transect the colony was collected from in “transect.position” with colonies with adjacent integers on the same transect at the same site being 10 meters apart, and whether the sample is included in the photographic, whole genome sequencing, or the raw proteomic and metabolomic datasets. The symbol “–” is used to indicate whether a sample was not collected on transect or not present in a given dataset.

| Genus | Name | ID | Date | Site | Transect | Photo | WGS | Prot | Met |
| --- | --- | --- | --- | --- | --- | --- | --- | --- | --- |
| Montipora | M1 | M001 | 230615 | Falalop Men's House | – | Yes | Yes | Yes | Yes |
| Montipora | M2 | M002 | 230615 | Falalop Men's House | – | Yes | Yes | Yes | Yes |
| Montipora | M3 | M003 | 230615 | Falalop Men's House | – | Yes | Yes | Yes | Yes |
| Montipora | M4 | M004 | 230615 | Falalop Men's House | – | Yes | Yes | Yes | Yes |
| Montipora | M5 | M005 | 230615 | Falalop Men's House | – | Yes | Yes | Yes | Yes |
| Montipora | M6 | M006 | 230615 | Falalop Men's House | – | Yes | Yes | Yes | Yes |
| Montipora | M7 | M007 | 230615 | Falalop Men's House | – | Yes | Yes | Yes | Yes |
| Montipora | M8 | M008 | 230615 | Falalop Men's House | – | Yes | Yes | Yes | Yes |
| Montipora | M9 | M009 | 230615 | Falalop Men's House | – | Yes | Yes | Yes | Yes |
| Montipora | M10 | M010 | 230615 | Falalop Men's House | – | Yes | Yes | Yes | Yes |
| Montipora | M11 | M011 | 230616 | Asor Landing | 1.1 | Yes | Yes | Yes | Yes |
| Montipora | M12 | M012 | 230616 | Asor Landing | 1.2 | Yes | Yes | Yes | Yes |
| Montipora | M13 | M013 | 230616 | Asor Landing | 1.3 | Yes | Yes | Yes | Yes |
| Montipora | M14 | M014 | 230616 | Asor Landing | 1.4 | Yes | Yes | Yes | Yes |
| Montipora | M15 | M015 | 230616 | Asor Landing | 1.5 | Yes | Yes | Yes | Yes |
| Montipora | M16 | M016 | 230616 | Asor Landing | 2.1 | Yes | Yes | Yes | Yes |
| Montipora | M17 | M017 | 230616 | Asor Landing | 2.2 | Yes | Yes | Yes | Yes |
| Montipora | M18 | M018 | 230616 | Asor Landing | 2.3 | Yes | Yes | Yes | Yes |
| Montipora | M19 | M019 | 230616 | Asor Landing | 2.4 | Yes | Yes | Yes | Yes |
| Montipora | M20 | M020 | 230616 | Asor Landing | 2.5 | Yes | Yes | Yes | Yes |
| Montipora | M21 | M021 | 230617 | MogMog Landing | 1.1 | Yes | Yes | Yes | Yes |
| Montipora | M22 | M022 | 230617 | MogMog Landing | 1.2 | Yes | Yes | Yes | Yes |
| Montipora | M23 | M023 | 230617 | MogMog Landing | 1.3 | Yes | Yes | Yes | Yes |
| Montipora | M24 | M024 | 230617 | MogMog Landing | 1.4 | Yes | Yes | Yes | Yes |
| Montipora | M25 | M025 | 230617 | MogMog Landing | 1.5 | Yes | Yes | Yes | Yes |
| Montipora | M26 | M026 | 230617 | MogMog Landing | 2.1 | Yes | Yes | Yes | Yes |
| Montipora | M27 | M027 | 230617 | MogMog Landing | 2.2 | Yes | Yes | Yes | Yes |
| Montipora | M28 | M028 | 230617 | MogMog Landing | 2.3 | Yes | Yes | Yes | Yes |
| Montipora | M29 | M029 | 230617 | MogMog Landing | 2.4 | Yes | Yes | – | Yes |
| Montipora | M30 | M030 | 230617 | MogMog Landing | 2.5 | Yes | Yes | Yes | Yes |
| Montipora | M31 | M031 | 230621 | Sorghlei | 1.1 | Yes | Yes | Yes | Yes |
| Montipora | M32 | M032 | 230621 | Sorghlei | 1.2 | Yes | Yes | Yes | Yes |
| Montipora | M33 | M033 | 230621 | Sorghlei | 1.3 | Yes | Yes | Yes | Yes |
| Montipora | M34 | M034 | 230621 | Sorghlei | 1.4 | Yes | Yes | Yes | Yes |
| Montipora | M35 | M035 | 230621 | Sorghlei | 1.5 | Yes | Yes | Yes | Yes |
| Montipora | M36 | M036 | 230621 | Sorghlei | 2.1 | Yes | Yes | Yes | Yes |
| Montipora | M37 | M037 | 230621 | Sorghlei | 2.2 | Yes | Yes | Yes | Yes |
| Montipora | M38 | M038 | 230621 | Sorghlei | 2.3 | Yes | Yes | Yes | Yes |
| Montipora | M39 | M039 | 230621 | Sorghlei | 2.4 | Yes | Yes | Yes | Yes |
| Montipora | M40 | M040 | 230621 | Sorghlei | 2.5 | Yes | Yes | Yes | Yes |
| Montipora | M41 | M041 | 230622 | Federali Landing | 1.1 | Yes | Yes | Yes | Yes |
| Montipora | M42 | M042 | 230622 | Federali Landing | 1.2 | Yes | Yes | Yes | Yes |
| Montipora | M43 | M043 | 230622 | Federali Landing | 1.3 | Yes | Yes | Yes | Yes |
| Montipora | M44 | M044 | 230622 | Federali Landing | 1.4 | Yes | Yes | Yes | Yes |
| Montipora | M45 | M045 | 230622 | Federali Landing | 1.5 | Yes | Yes | Yes | Yes |
| Montipora | M46 | M046 | 230622 | Federali Landing | 2.1 | Yes | Yes | Yes | Yes |
| Montipora | M47 | M047 | 230622 | Federali Landing | 2.2 | Yes | Yes | Yes | Yes |
| Montipora | M48 | M048 | 230622 | Federali Landing | 2.3 | Yes | Yes | Yes | Yes |
| Montipora | M49 | M049 | 230622 | Federali Landing | 2.4 | Yes | Yes | Yes | Yes |
| Montipora | M50 | M050 | 230622 | Federali Landing | 2.5 | Yes | Yes | Yes | Yes |
| Montipora | M51 | M051 | 230623 | Fetabul | 1.1 | Yes | Yes | Yes | Yes |
| Montipora | M52 | M052 | 230623 | Fetabul | 1.2 | Yes | Yes | Yes | Yes |
| Montipora | M53 | M053 | 230623 | Fetabul | 1.3 | Yes | Yes | Yes | Yes |
| Montipora | M54 | M054 | 230623 | Fetabul | 1.4 | Yes | Yes | Yes | Yes |
| Montipora | M55 | M055 | 230623 | Fetabul | 1.5 | Yes | Yes | Yes | Yes |
| Montipora | M56 | M056 | 230623 | Fetabul | 2.1 | Yes | Yes | Yes | Yes |
| Montipora | M57 | M057 | 230623 | Fetabul | 2.2 | Yes | Yes | Yes | Yes |
| Montipora | M58 | M058 | 230623 | Fetabul | 2.3 | Yes | Yes | Yes | Yes |
| Montipora | M59 | M059 | 230623 | Fetabul | 2.4 | Yes | Yes | Yes | Yes |
| Montipora | M60 | M060 | 230623 | Fetabul | 2.5 | Yes | Yes | Yes | Yes |
| Montipora | M71 | M071 | 230624 | Pigleilei Inside | 1.1 | Yes | Yes | Yes | Yes |
| Montipora | M72 | M072 | 230624 | Pigleilei Inside | 1.2 | Yes | Yes | Yes | Yes |
| Montipora | M73 | M073 | 230624 | Pigleilei Inside | 1.3 | Yes | Yes | Yes | Yes |
| Montipora | M74 | M074 | 230624 | Pigleilei Inside | 1.4 | Yes | Yes | Yes | Yes |
| Montipora | M75 | M075 | 230624 | Pigleilei Inside | 1.5 | Yes | Yes | Yes | Yes |
| Montipora | M76 | M076 | 230624 | Pigleilei Inside | 2.1 | Yes | Yes | Yes | Yes |
| Montipora | M77 | M077 | 230624 | Pigleilei Inside | 2.2 | Yes | Yes | Yes | Yes |
| Montipora | M78 | M078 | 230624 | Pigleilei Inside | 2.3 | Yes | Yes | Yes | Yes |
| Montipora | M79 | M079 | 230624 | Pigleilei Inside | 2.4 | Yes | Yes | Yes | Yes |
| Montipora | M80 | M080 | 230624 | Pigleilei Inside | 2.5 | Yes | Yes | Yes | Yes |
| Montipora | M81 | M081 | 230624 | Laam | 1.1 | Yes | Yes | Yes | Yes |
| Montipora | M82 | M082 | 230624 | Laam | 1.2 | Yes | Yes | Yes | Yes |
| Montipora | M83 | M083 | 230624 | Laam | 1.3 | Yes | Yes | Yes | Yes |
| Montipora | M84 | M084 | 230624 | Laam | 1.4 | Yes | Yes | Yes | Yes |
| Montipora | M85 | M085 | 230624 | Laam | 1.5 | Yes | Yes | Yes | Yes |
| Montipora | M86 | M086 | 230624 | Laam | 2.1 | Yes | Yes | Yes | Yes |
| Montipora | M87 | M087 | 230624 | Laam | 2.2 | Yes | Yes | Yes | Yes |
| Montipora | M88 | M088 | 230624 | Laam | 2.3 | Yes | Yes | Yes | Yes |
| Montipora | M89 | M089 | 230624 | Laam | 2.4 | Yes | Yes | Yes | Yes |
| Montipora | M90 | M090 | 230624 | Laam | 2.5 | Yes | Yes | Yes | Yes |
| Montipora | M100 | M091 | 230625 | Sohng | 1.1 | Yes | Yes | Yes | Yes |
| Montipora | M91 | M092 | 230625 | Sohng | 1.2 | Yes | Yes | Yes | Yes |
| Montipora | M92 | M093 | 230625 | Sohng | 1.3 | Yes | Yes | Yes | Yes |
| Montipora | M93 | M094 | 230625 | Sohng | 1.4 | Yes | Yes | Yes | Yes |
| Montipora | M94 | M095 | 230625 | Sohng | 1.5 | Yes | Yes | Yes | Yes |
| Montipora | M95 | M096 | 230625 | Sohng | 2.1 | Yes | Yes | Yes | Yes |
| Montipora | M96 | M097 | 230625 | Sohng | 2.2 | Yes | Yes | Yes | Yes |
| Montipora | M97 | M098 | 230625 | Sohng | 2.3 | Yes | Yes | Yes | Yes |
| Montipora | M98 | M099 | 230625 | Sohng | 2.4 | Yes | Yes | Yes | Yes |
| Montipora | M99 | M100 | 230625 | Sohng | 2.5 | Yes | Yes | Yes | Yes |
| Montipora | M101 | M101 | 230626 | Yealeil Inside | 1.1 | Yes | Yes | Yes | Yes |
| Montipora | M102 | M102 | 230626 | Yealeil Inside | 1.2 | Yes | Yes | Yes | Yes |
| Montipora | M103 | M103 | 230626 | Yealeil Inside | 1.3 | Yes | Yes | Yes | Yes |
| Montipora | M104 | M104 | 230626 | Yealeil Inside | 1.4 | Yes | Yes | Yes | Yes |
| Montipora | M105 | M105 | 230626 | Yealeil Inside | 1.5 | Yes | Yes | Yes | Yes |
| Montipora | M106 | M106 | 230626 | Yealeil Inside | 2.1 | Yes | Yes | Yes | Yes |
| Montipora | M107 | M107 | 230626 | Yealeil Inside | 2.2 | Yes | Yes | Yes | Yes |
| Montipora | M108 | M108 | 230626 | Yealeil Inside | 2.3 | Yes | Yes | Yes | Yes |
| Montipora | M109 | M109 | 230626 | Yealeil Inside | 2.4 | Yes | Yes | Yes | Yes |
| Montipora | M110 | M110 | 230626 | Yealeil Inside | 2.5 | Yes | Yes | Yes | Yes |

**Table 3.** Summary of *Pocillopora spp.* samples collected. Columns from left to right correspond to: genus, sample id, collection date, site name, the position on the transect the colony was collected from in “transect.position” with colonies with adjacent integers on the same transect at the same site being 10 meters apart, and whether the sample is included in the photographic dataset or the proteomic and metabolomic datasets. The symbol “–” is used to indicate whether a sample was not collected on transect or not present in a given dataset.

| Genus | Sample | Date | Site | Transect | Photo | Prot | Met |
| --- | --- | --- | --- | --- | --- | --- | --- |
| Pocillopora | M66 | 230618 | BulBul Outside | - | Yes | Yes | Yes |
| Pocillopora | M67 | 230618 | BulBul Outside | - | Yes | Yes | Yes |
| Pocillopora | M68 | 230618 | BulBul Outside | - | Yes | Yes | Yes |
| Pocillopora | M69 | 230618 | BulBul Outside | - | Yes | Yes | Yes |
| Pocillopora | M70 | 230618 | BulBul Outside | - | Yes | Yes | Yes |
| Pocillopora | P11 | 230615 | Falalop Men's House | - | Yes | Yes | Yes |
| Pocillopora | P12 | 230615 | Falalop Men's House | - | Yes | Yes | Yes |
| Pocillopora | P13 | 230615 | Falalop Men's House | - | Yes | Yes | Yes |
| Pocillopora | P14 | 230616 | Falalop Men's House | - | Yes | Yes | Yes |
| Pocillopora | P15 | 230616 | Falalop Men's House | - | Yes | Yes | Yes |
| Pocillopora | P16 | 230616 | Falalop Men's House | - | Yes | Yes | Yes |
| Pocillopora | P17 | 230616 | Falalop Men's House | - | Yes | Yes | Yes |
| Pocillopora | P18 | 230616 | Falalop Men's House | - | Yes | Yes | Yes |
| Pocillopora | P19 | 230616 | Falalop Men's House | - | Yes | Yes | Yes |
| Pocillopora | P20 | 230616 | Falalop Men's House | - | Yes | Yes | Yes |
| Pocillopora | P201 | 230617 | MogMog Landing | 1.1 | Yes | Yes | Yes |
| Pocillopora | P202 | 230617 | MogMog Landing | 1.2 | Yes | Yes | Yes |
| Pocillopora | P203 | 230617 | MogMog Landing | 1.3 | Yes | Yes | Yes |
| Pocillopora | P204 | 230617 | MogMog Landing | 1.4 | Yes | Yes | Yes |
| Pocillopora | P205 | 230617 | MogMog Landing | 1.5 | Yes | Yes | Yes |
| Pocillopora | P206 | 230617 | MogMog Landing | 2.1 | Yes | Yes | Yes |
| Pocillopora | P207 | 230617 | MogMog Landing | 2.2 | Yes | Yes | Yes |
| Pocillopora | P208 | 230617 | MogMog Landing | 2.3 | Yes | Yes | Yes |
| Pocillopora | P209 | 230617 | MogMog Landing | 2.4 | Yes | Yes | Yes |
| Pocillopora | P210 | 230617 | MogMog Landing | 2.5 | Yes | Yes | Yes |

**Table 4.** Summary of *Acropora spp.* samples collected. Columns from left to right correspond to: genus, sample id, collection date, site name, the position on the transect the colony was collected from in “transect.position” with colonies with adjacent integers on the same transect at the same site being 10 meters apart, and whether the sample is included in the photographic dataset or the proteomic and metabolomic datasets. The symbol “–” is used to indicate whether a sample was not collected on transect or not present in a given dataset.

| Genus | Sample | Date | Site | Transect | Photo | Prot | Met |
| --- | --- | --- | --- | --- | --- | --- | --- |
| Acropora | A21 | 230615 | Falalop Men's House | - | Yes | Yes | Yes |
| Acropora | A22 | 230615 | Falalop Men's House | - | Yes | Yes | Yes |
| Acropora | A23 | 230615 | Falalop Men's House | - | Yes | Yes | Yes |
| Acropora | A24 | 230615 | Falalop Men's House | - | Yes | Yes | Yes |
| Acropora | A25 | 230615 | Falalop Men's House | - | Yes | Yes | Yes |
| Acropora | A26 | 230615 | Falalop Men's House | - | Yes | Yes | Yes |
| Acropora | A27 | 230615 | Falalop Men's House | - | Yes | Yes | - |
| Acropora | A28 | 230615 | Falalop Men's House | - | Yes | Yes | - |
| Acropora | A29 | 230615 | Falalop Men's House | - | Yes | Yes | - |
| Acropora | A30 | 230615 | Falalop Men's House | - | Yes | Yes | - |
| Acropora | A306 | 230617 | MogMog Landing | 1.1 | - | - | Yes |
| Acropora | A309 | 230617 | MogMog Landing | 1.2 | - | Yes | Yes |
| Acropora | A315 | 230617 | MogMog Landing | 1.3 | - | Yes | Yes |
| Acropora | A323 | 230617 | MogMog Landing | 1.4 | Yes | Yes | Yes |
| Acropora | A325 | 230617 | MogMog Landing | 1.5 | Yes | Yes | Yes |
| Acropora | A326 | 230617 | MogMog Landing | 2.1 | Yes | Yes | Yes |
| Acropora | A328 | 230617 | MogMog Landing | 2.2 | Yes | Yes | Yes |
| Acropora | A331 | 230617 | MogMog Landing | 2.3 | - | Yes | Yes |
| Acropora | A338 | 230617 | MogMog Landing | - | - | Yes | - |
| Acropora | A339 | 230617 | MogMog Landing | 2.4 | - | Yes | Yes |
| Acropora | A340 | 230617 | MogMog Landing | 2.5 | - | Yes | Yes |
| Acropora | M61 | 230618 | BulBul Outside | - | Yes | Yes | Yes |
| Acropora | M62 | 230618 | BulBul Outside | - | Yes | Yes | Yes |
| Acropora | M63 | 230618 | BulBul Outside | - | Yes | Yes | Yes |
| Acropora | M64 | 230618 | BulBul Outside | - | Yes | Yes | Yes |
| Acropora | M65 | 230618 | BulBul Outside | - | Yes | Yes | Yes |

### Whole Genome Sequencing (WGS) Dataset

#### DNA extraction and sequencing

DNA was extracted using the Zymo Quick MiniPrep Plus Kit, following the kit’s instructions for samples in DNA/RNA Shield and eluted in 100 µL of DNase/RNase-free water (**Table S1**). DNA was quantified with a ThermoFisher Qubit 2.0 fluorometer and integrity was determined by running 10 µL aliquots for each sample on a 1% agarose gel in Tris Acetate-EDTA buffer (100 V). Extracted gDNA was sent for library preparation and sequencing at Azenta Life Sciences (South Plainfield, NJ). Library preparation and sequencing followed the Azenta protocol for Short Read Non-Human Whole Genome Sequencing, targeting a read length of 250 bp and ∼30x depth of coverage for a 1,000 Mbp genome (30 Gbp total sequenced bases). This depth of coverage was chosen given that the genome size of *Montipora capitata* is estimated to be around 800 Mbp [15] and that reads would also be generated from the algal symbionts and prokaryotic microbiome, 1,000 Mbp was used as a lenient overestimate to ensure that sufficient coverage of the coral host was recovered for downstream analyses.

#### Read processing and variant calling

The read processing workflow used in this study was adapted from the DeepVariant workflow [16]. First, raw reads were processed using fastp v0.23.2 [17] with default parameters. Both raw and trimmed reads were checked for quality using fastqc v0.12.1 [18] and multiqc v1.14 (default parameters) [19]. Trimmed reads were then aligned to the *Montipora sp.1 aff. capitata* reference genome [13] using bwa-mem v2.2.1 [20], marking shorter split hits as secondary (-M) and flagging all unpaired paired-end reads as secondary alignments (-a). Mapped reads were then coordinate-sorted and quality filtered using Samtools v1.13 [21], retaining only primary alignments (-F 0x100) with MAPQ (Phred-scale) scores greater than 20 (-q 20). Samtools and Qualimap v2.2.2a [22] were then used to generate statistics for quality control metrics for both raw and clean mapped reads including duplicates, number of primary and secondary alignments, depth, and GC content and insert size of aligned reads. Finally, variants were called from each of the aligned BAM files using DeepVariant v1.4.0 with the WGS model (--model_type=WGS), outputting individual gVCFs (--output_gvcf) and using a gVCF bin size of 1 (--make_examples_extra_args gvcf_gq_binsize=1) [16]. Accurate inferences for several traditional metrics of population divergence and relatedness (e.g., nucleotide diversity (π), nucleotide divergence (D_XY_), and Tajima’s D statistic) require data on both variant and invariant sites [23]. Because the DeepVariant cohort-calling method does not include an option for generating an AllSites VCF, we used the GATK joint genotyping method [24] (gatk GenotypeGVCFs) to generate a population-level AllSites VCF (-all-sites) using the gVCFs from DeepVariant. Lastly, we filtered the resulting population-level VCF for depth, missingness, minor allele frequency, and genotype quality. Invariant and variant sites were filtered separately and then recombined.

The following parameters were used to filter variant-only sites (Genotype Quality [minQ] > 30, removal of indels [remove-indels], minor allele frequency [maf] > 0.05). Depth and missingness filters were then applied independently to both invariant and variant site VCFs (minimum genotype depth [minDP] 10, maximum genotype depth [maxDP] 70, minimum mean site depth [min-meanDP] 10, maximum mean site depth [max-meanDP] 70, site missingness < 0.2 [max-missing 0.8]) prior to recombining the files for QC and downstream analyses. VCFtools was used to generate statistics, including genotype quality, depth, missingness, and inbreeding coefficients, for quality control before and after filtering. We have made both the unfiltered and filtered population-level AllSites VCFs as well as the filtered variant-only population-level VCF available for public use, as well as the raw sequencing reads. See **Data Availability** for further details.

#### Symbiont community profiling

We inferred the major symbiont clades present in the *Montipora* sp. samples using Kraken, a *k*-mer-based tool for taxonomic classification of metagenomic reads, using a workflow for identification of symbionts from coral holobiont WGS data described by Zhang et al. 2022 [25]. We constructed a custom Kraken database using two *Montipora* reference genomes, six Symbiodiniaceae reference genomes representing the most common coral associated genera, and the built-in Kraken bacteria nucleotide library (**Table S2**). Summaries of sample composition by taxonomic group in the database were generated using the Kraken2 MPA-style report (kraken2 --use-mpa-style) function.

### Proteomic Dataset

#### Protein purification

Proteins were extracted using a Radio-Immunoprecipitation Assay (RIPA) lysis buffer, composed of Tris-HCl (50 mM), NaCl (150 mM), SDS (1%), and DI water. The buffer was chilled on ice until use; one cOmplete mini EDTA-free tablet was dissolved in 10 mL of lysis buffer immediately before extraction. A total of 500 μL of ice-cold RIPA lysis buffer was added to a tube filled with 0.5 mm silica beads. To extract total proteins preserved in RNAlater, nubbins were removed from AllProtect solution and rinsed in 1xPBS. The frozen nubbins were then immediately placed in the RIPA buffer lysed for 5 minutes using a vortex mixer set to maximum speed. The samples were then incubated on ice for 30 minutes, before centrifuging (at 10,000 rcf at 4°C). The supernatants are clarified by a second centrifugation (10,000 rcf for 5 minutes). After extraction, aliquots of protein extracts are taken for determination of protein concentration using the BCA method, and the remaining extracts stored at −80°C for downstream applications.

#### Mass Spectrometry and Data Analysis

To prepare crude protein extracts for mass spectrometry, 20 μg of total protein were aliquoted from each sample and diluted to 100 µL in reconstitute buffer (50 mM HEPEs, pH8, 1% SDS and 50 mM EDTA). Then, samples were incubated in 5 mM DTT for 30 min at 60°C and subjected to free cysteine alkylation with 20 mM iodoacetamide for 1 hour at room temperature in dark conditions. The samples were then further digested with SP3 beads digestion [26] with trypsin (sequencing grade, Thermo Scientific Cat#90058) in 100 mM ammonium bicarbonate, 2 mM CaCl2 and incubated at 37°C for 1 hour. After digestion, samples were concentrated under vacuum to 20 µL and then diluted to 100 µL in 5% formic acid 60% acetonitrile. Samples were incubated at room temp for 30 min and then insoluble contaminants were precipitated by centrifugation at 25,000 rcf for 10 min. Clarified supernatants were dried under vacuum to about 10 µL and then desalted using Stage-Tip method prior to analysis.

Samples were analyzed using Liquid Chromatography–Mass Spectrometry (LC-MS) using Nano LC-MS/MS (Dionex Ultimate 3000 RLSCnano System, Thermofisher) interfaced with Eclipse (Thermofisher). Samples (1 µg) were loaded on to a fused silica trap column Acclaim PepMap 100, 75 µm x 2 cm (ThermoFisher). After washing for 5 min at 5 µL/min with 0.1% TFA, the trap column was brought in-line with an analytical column (Nanoease MZ peptide BEH C18, 130A, 1.7 µm, 75 µm x 250 mm, Waters) for LC-MS/MS. Peptides were fractionated at 300 nL/min using a segmented linear gradient 4-15% B in 30min (where A: 0.2% formic acid, and B: 0.16% formic acid, 80% acetonitrile), 15-25% B in 40min, 25-50% B in 44min, and 50-90% B in 11min. Solution B then returns at 4% for 5 minutes for the next run.

A DIA (Data Independent Acquisition) workflow was used to analyze the eluted peptides. MS scan range was set at 400-1200, resolution 12,000 with AGC set at 3E6 and ion time set as auto. An 8 m/z window was set to sequentially isolate (AGC 4E5 and ion time set at auto) and fragment the ions in C-trap with relative collision energy of 30. The MSMS were recorded with Resolution of 30,000. Raw data were analyzed with predicted library from customer supplied databases for each coral genus using DIA NN 1.8.1 (recommended settings) [27]. The custom libraries were built to include predicted proteins from both the host (*Montipora capitata* [15] [*Montipora* sp.], *Acropora millepora* [28] [*Acropora spp*.], and *Pocillopora verrucosa* [29] [*Pocillopora* spp.]) and symbiont (with one representative each for the two the most common symbiont clades, *Cladocopoium goreaei* [30] and *Durusdinium trenchii* [31]) in each custom library.

#### Proteomic data cleaning

Proteomic data cleaning was conducted in RStudio v2024.04.2+764 using R v4.4.1 and the protti R package v0.9.0 [32]. This package was used to clean, normalize, and generate pre- and post-cleaning reports for each of the three coral genera independently.

Assessments of peak intensity distribution, data completeness, and principal components analysis (PCA) were performed before and after cleaning. To create a high-quality host protein group library, LC-MS results were filtered to exclude proteins identified as symbiont-derived. Proteins called with low confidence (e.g., with a posterior error probability [PEP] and a protein group Q-value less than 0.01, and precursor quantities below 1) were also removed. Protein intensities were log2-transformed using the dplyr *mutate* and base R *log2* functions. Outlier samples were identified by PCA using the protti *qc_pca* function. Samples that were visual outliers on PC1 and PC2 and that had low data completeness were removed (see **Results** for details). Missingness was then calculated for each site. Proteins were retained if they were missing in fewer than 40% of samples in any site (that is, a protein was retained if it was considered present at each of the 10 collection sites, with a protein considered present at a site if it was detected in >60% of samples collected from that site). This threshold was chosen because *missForest* has been benchmarked to be effective at imputing missing values with missingness of up to 30% [33]. For the remaining high-confidence observations, missing values were imputed using the *missForest* function [33] with default parameters (maxiter = 10, ntree = 100) and a set seed of 124. Finally, log_2_-transformed peak intensities were median-normalized per sample using the protti *normalise* function to account for variance in run intensity due to injection order and batch (**Table S3**), resulting in a final cleaned protein abundance matrix for each genus.

### Metabolomic Dataset

#### Metabolomic extraction and LC-MS

Metabolites were extracted using a buffer optimized for water-soluble polar metabolite analysis on LC-MS (40:40:20 [Methanol:Acetonitrile:Water] (v/v/v) + 0.1 M Formic Acid). Buffer was prepared in advance and stored at −20°C until usage. Immediately preceding the metabolite extraction, 1 mL of ice-cold lysis buffer was added to a tube filled with 0.5 mm silica beads. The frozen coral nubbins (∼0.2-0.4 g) were then immediately placed in lysis buffer and vortexed at maximum speed for 5 minutes to homogenize the coral tissue. Tissue homogenates were then transferred to a 1.5 mL conical safe-lock Eppendorf tube and centrifuged at 4°C at 15,000g for 10 minutes. The supernatant was transferred in increments of 125 µL to a clean 1.5 mL conical safe-lock Eppendorf tube, careful not to disturb the pellet. Exactly 11 µL of 15% NH_4_HCO_3_ was added to every 125 µL of supernatant transferred to neutralize the acid in the extraction buffer.

The sample was vortexed for 10 seconds and centrifuged at 4°C at 15,000g for 3 minutes. Following centrifugation, 500 µL of the supernatant was transferred to a second clean 1.5 mL conical safe-lock Eppendorf tube. This was the final metabolite extract and was stored at -80°C until sample processing for LC-MS.

Metabolite samples were shipped to the University of Connecticut Proteomics & Metabolomics Facility for Liquid Chromatography–Mass Spectrometry. Methanol extracts of the coral samples were diluted (1:3, v/v) in Fisher Optima LC-MS grade water in pre-weighed 5 mL plastic tubes (Eppendorf, Germany). The samples were then frozen, and lyophilized (Labconco, MO) to obtain the masses of the dried extract which were later reconstituted in acetonitrile: methanol: water (40:40:20, v/v/v) to a final concentration of 5 mg/mL for injection.

For metabolomic analysis, an aliquot of each sample was analyzed via untargeted ultra-high performance liquid chromatography coupled to tandem mass spectrometry using an Acquity UPLC and Synapt G2-Si mass spectrometer system (Waters Corp., MA). Hydrophilic interaction liquid chromatography separations were performed on an XBridge Premier BEH Amide column (150 x 2.1 mm, 2.5 µm particle size; Waters Corp., MA) using water:acetonitrile (95:5, v/v) with 20 mM acetic acid and 40 mM ammonium hydroxide (Solvent A) and water: acetonitrile (20:80, v/v) with 20 mM acetic acid and 40 mM ammonium hydroxide (Solvent B). The mobile phase gradient was as follows: 0 - 3 min: 100% B; 3.2 - 6.2 min: 90% B; 6.5 - 10.5 min: 80% B; 10.7 - 13.5 min: 70% B; 13.7 - 16 min: 45% B; and 16.5 - 22 min: 100% B. The flow rate was 300 µL/min and the sample injection volume was 10 µL. The column temperature was maintained at 25°C and the autosampler was set to 4°C. The analysis was performed using electrospray ionization in both positive and negative modes. The scan range for MS and MSe acquisitions were 72 to 1000 Da and 40 to 1000 Da, respectively. The acquisitions were made in the resolution acquisition mode. A collision energy ramp of 5 to 40 V was applied for MSe acquisition. Leucine enkephalin at 400 pg/µL, at an infusion flow rate of 10 µL/min, was used for real-time lockspray correction. The capillary voltage was set to 2.5 kV and the sampling cone at 40 V. The desolvation temperature was set to 250 °C, cone gas flow set to 50 L/hr, and the desolvation gas flow set to 600 L/hr.

Raw data files from positive and negative ionization modes were separately exported to Progenesis QI (v2.4) software for data processing, which included cross-sample retention time alignment, peak picking, deconvolution, normalization, and metabolite annotation. METLIN MS/MS library was used as a compound identification database [34]. The search parameters were set at 10 ppm precursor tolerance and 20 ppm fragment tolerance. This was done separately for positive mode and negative mode ionization.

#### Metabolomic data cleaning

Metabolomic data cleaning was conducted on positive and negative ionization peak intensity tables separately for each genus in RStudio (v2024.04.2+764) using R (v4.4.1). The TidyMass (v1.0.9) package data cleaning workflow (https://www.tidymass.org/docs/chapter6/1-data_cleaning/) was used to filter out noisy metabolites, detect and remove outlier metabolites and samples, impute missing values, normalize across samples, log_e_ scale the raw abundance values, and generate pre- and post-cleaning reports for the data [35]. Assessments of intensity distribution across samples, data completeness, relative standard deviation across metabolites, and PCA were performed before and after cleaning (**Figs. 9-11**). Noisy metabolites were filtered out following the using the threshold *MV.lim* = 0.4, that is, metabolites with a missingness frequency of 40% or higher across all samples were removed [33]. This threshold allows for effective missing value imputation using *missForest*, which has been benchmarked using missingness thresholds of 30% or lower [33]. Following filtering, the *detect_outlier* R function was used to identify and remove outliers. Samples were considered outliers if they met the requirements for at least two of the four *detect_outlier* tests (*according_to_na, according_to_pc_sd, according_to_pc_mad*, and *according_to_distance*). Missing values were imputed and replaced using the random forest method implemented in the *impute_mv* R function. Data was normalized using the “median” normalization method, implemented in the *normalize_data* R function. Data was integrated using the “subject_median” method from the *integrate_data* R function. The abundance matrices were log-scaled using the base R *log()* function. Finally, variance due to injection order was removed using the limma v3.58.1 R package [38]. We performed principal components analysis (PCA) before and after to confirm variance was removed from the data using base R *prcomp* function and R package ggplot2 (**Table S4**; **Fig. 9C-11C**).

### Quality Control

#### WGS dataset

##### Read processing and host mapping

High quality genomic data was generated from Illumina paired-end sequencing for all 100 *Montipora* sp. colonies sampled. On average, 204 M ± 9.31 M (mean ± standard deviation [SD]) raw PE reads were generated per sample (**Table S5**), with 202 M ± 1.84 M reads remaining after filtering (**Table S6**). MultiQC summaries before and after quality trimming showed that all samples displayed “Good” Phred scores (30 or higher), consistent GC content, low duplication levels both before and after filtering (**Fig. 2**). MultiQC plots after quality trimming indicate that adaptors had been effectively removed (**Fig. 2G**). After removing non-primary reads and reads with a mapping score (Phred scale) less than 20, we observed high mapping rates 95.86% ± 0.28% to the Ulithi *Montipora* sp. 1 *aff. capitata* genome with low error rates (2.08% ± 0.05%) and high mean coverage (35.19× ± 3.25×). On average, 88.56% ± 0.82% mapped reads had a coverage greater than 10×. A minimum of 128.40 M reads (153 M ± 1.28 M) were used for variant calling on each sample (**Table S8**).

**Figure 2.**
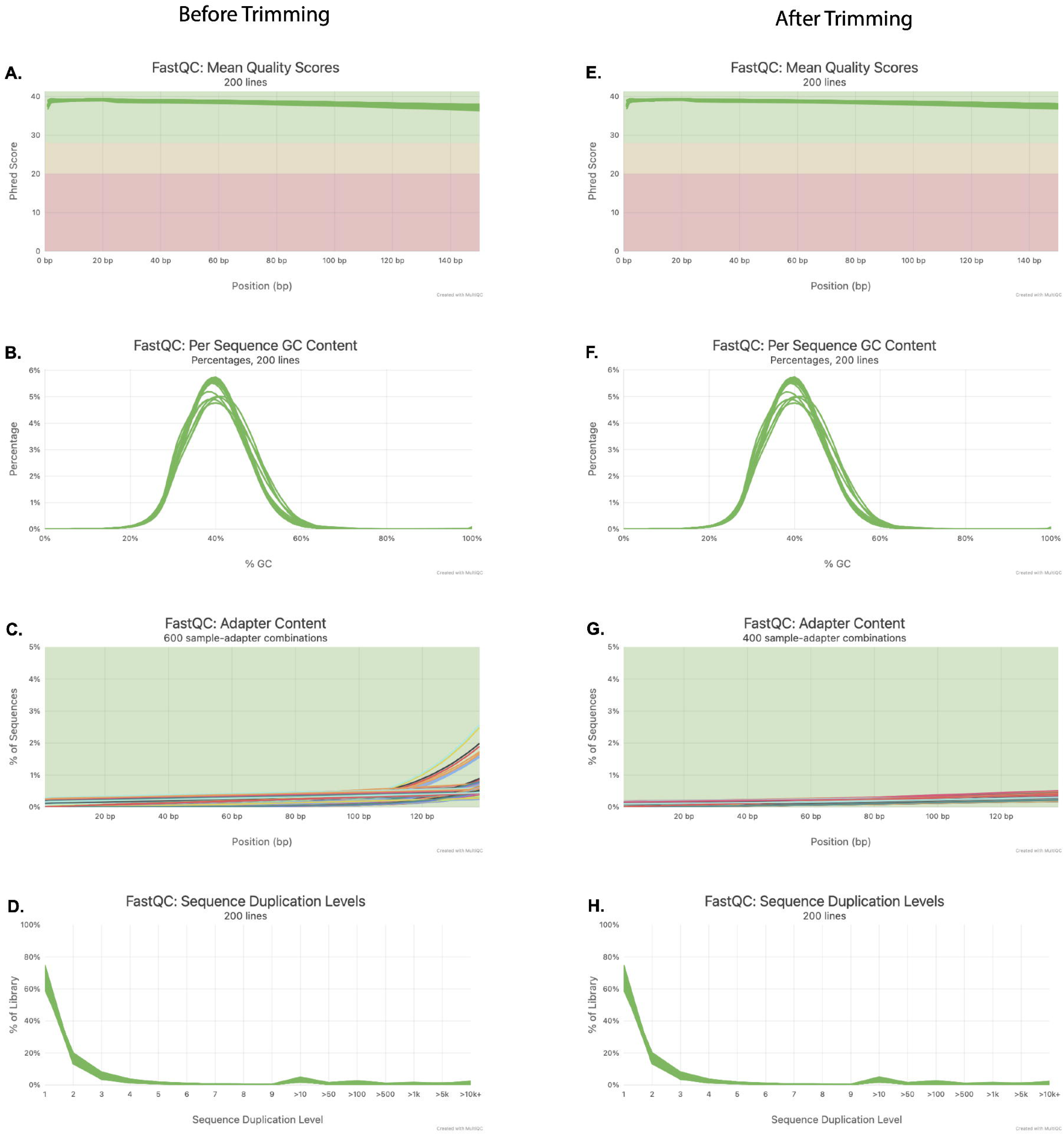
Quality control of *Montipora* WGS data. (A-D) Raw sequencing data. **(E-H)** Quality filtered sequencing data. **(A, E)** Mean quality value across each base position in the read. **(B, F)** Distribution of GC content. **(C, G)** Percentage of reads containing an Illumina adapter sequence across each base position in the read. **(D, H)** The distribution of sequence duplication levels across the reads in the library. In plots **(A, E, C, G)** background color indicates sequence quality (green, very good; yellow, reasonable; red, poor). In **(A-B, D-F, H)** green lines correspond to files that passed the FastQC quality control filters.

##### Variant calling

We used all 100 *Montipora* sp. colonies to call variants using the joint DeepVariant-GATK approach, obtaining an unfiltered AllSites VCF containing 514,240,230 sites with 24,852,151 variants, including 18,497,454 Single Nucleotide Polymorphisms (SNPs; 18,342,676 [99.16%] biallelic) and 2,332,768 indels. A total of 331,409,627 sites remained after filtering, with 428,727 biallelic SNPs that passed quality filtering parameters (genotype quality scores >30 [**Figs. 4A, D**], mean depth per individual 10× - 60× [**Figs. 4B, E**], and missingness <0.25 [**Figs. 4C, F**]). All individual samples had acceptable mean site depth (10× - 60×; **Figs. 3A, D**) and average missingness (<0.25; **Figs. 3B, E**). Mean depth, inbreeding coefficients, and missingness shifted for each individual after filtering likely due to removal of a small proportion of sites with extremely high coverage and removal of genotype calls with depths outside of the acceptable range (**Fig. 3**).

**Figure 3.**
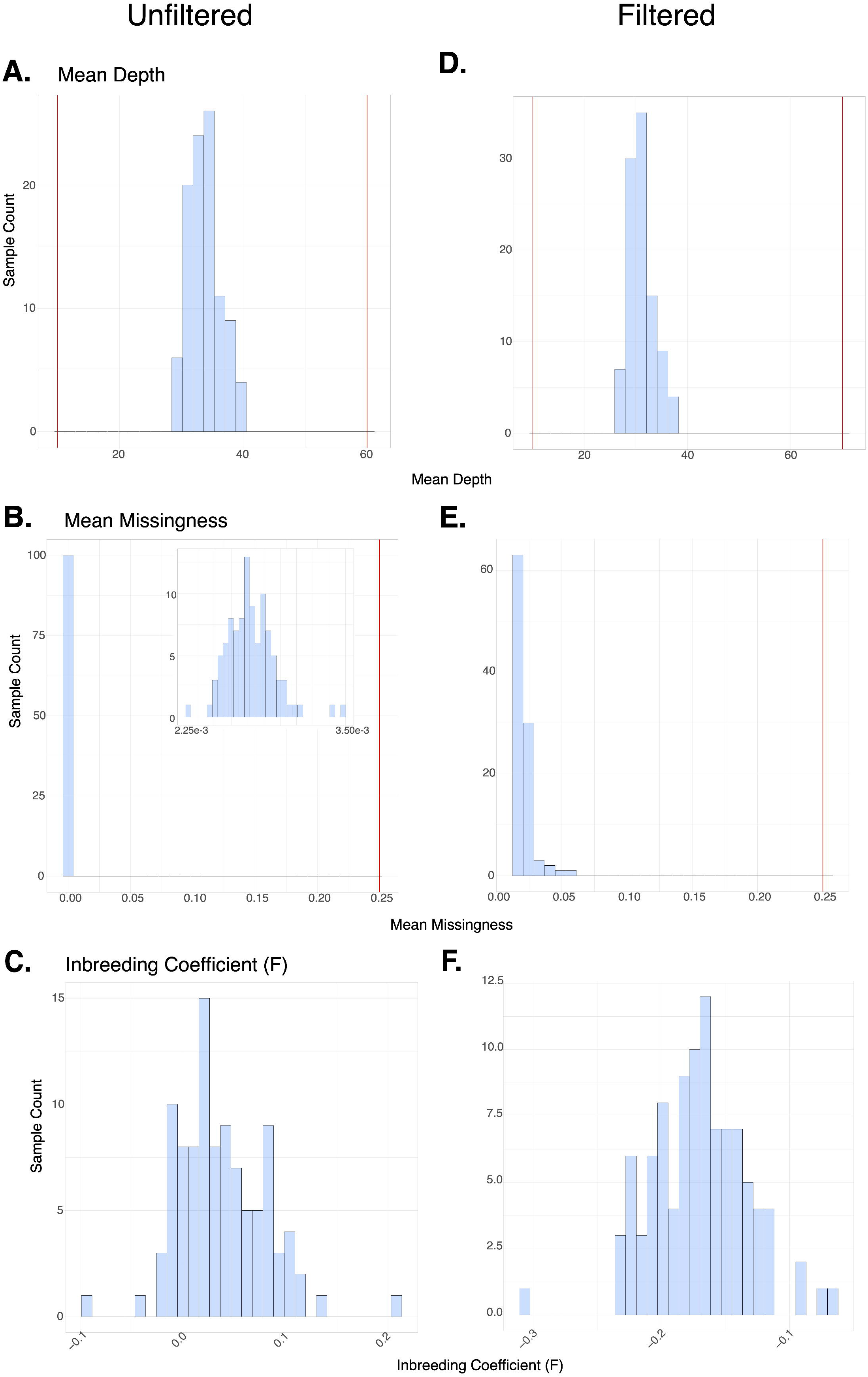
Individual-level quality control of *Montipora* variant calls. (A-C) Metrics calculated before quality filtering. **(D-F)** Metrics calculated after quality filtering. **(A, D)** Distribution of mean sequencing depth per sample. **(B, E)** Distribution of the average fraction of missing genotypes per sample. **(C, F)** Distribution of per-individual heterozygosity, reported as the inbreeding coefficient (*F*), calculated from biallelic SNPs. Red vertical lines indicate filtering thresholds applied during variant quality control.

**Figure 4.**
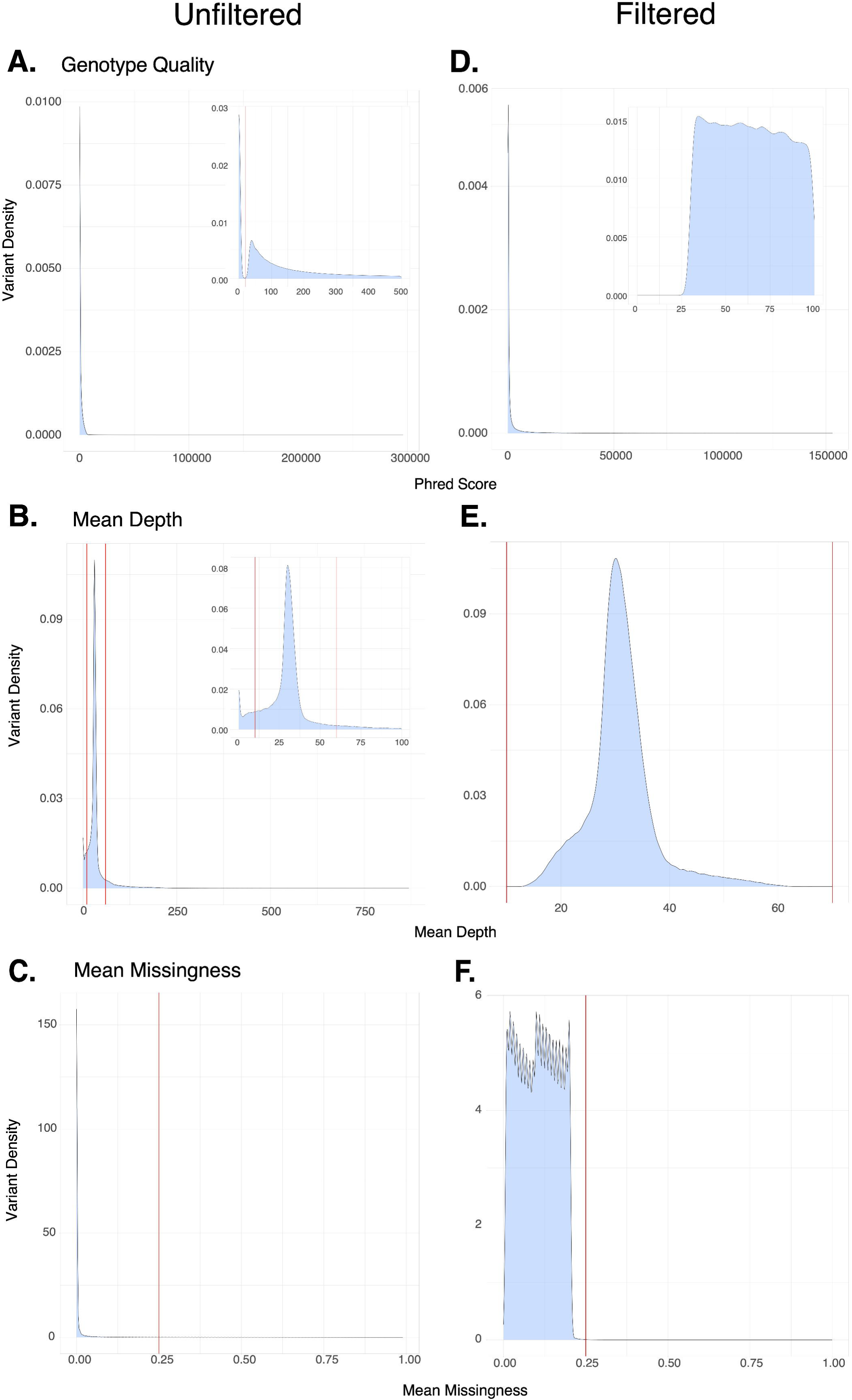
Site-level quality control of *Montipora* variant calls. (A-C) Metrics calculated before quality filtering. **(D-F)** Metrics calculated after quality filtering. **(A, D)** Distribution of genotype quality (Phred-scaled score) across variant sites. **(B, E)** Distribution of mean sequencing depth per site. **(C, F)** Distribution of mean genotype missingness per site. Red vertical lines indicate filtering thresholds applied during variant quality control.

##### Symbiont Composition

Kraken taxonomically classified a minimum of 81.21 M reads per sample (98.35 M ± 0.89 M [mean ± SD]) reads to either *Montipora* (host), Symbiodiniaceae, or bacteria given our custom-built database, which included two *Montipora* genomes, six Symbiodiniaceae genomes, and the Kraken built-in bacteria library (**Table S2**). Most reads were classified as host (98.33% ± 0.61%) with 0.57 - 3.83% (1.51% ± 0.58%) classified as symbiont, and less than 1% classified as bacterial (0.16% ± 0.14%). The dominant symbiont genus for all samples was *Cladocopium* (0.79% ± 0.05%; **Fig. 5**), with *Fugacium* (0.07% ± 0.02%) and *Symbiodinium* (0.05% ± 0.02%) present in low amounts (**Table S9**). There were no stark differences in the relative proportions of symbiont genera across the sites (**Fig. 5**). However, three samples from Fetabul – M54, M59, and M60 – had a marginally higher proportion of *Symbiodinium* (0.21, 0.15, 0.12, respectively) and a correspondingly lower proportion of *Cladocopium* (0.56, 0.60, and 0.69, respectively) relative to the average.

**Figure 5.**
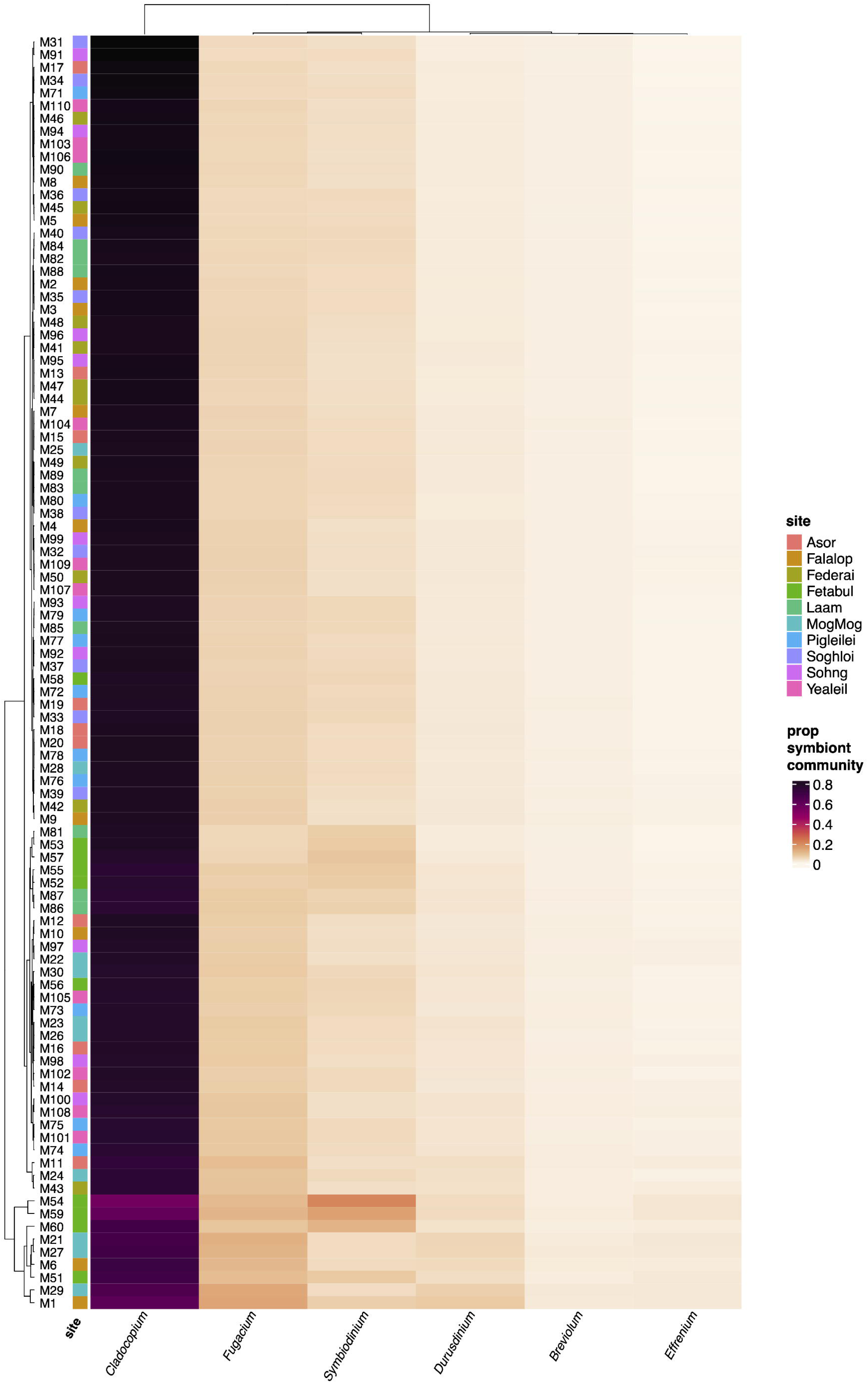
Symbiont community composition across *Montipora* samples. Heatmap showing the relative abundance of major Symbiodiniaceae genera across all samples. Columns represent symbiont genera and rows represent individual coral samples. Cell color indicates the proportion of Symbiodiniaceae reads from each genus. Samples are hierarchically clustered based on the proportion of reads from each genus. The colored bar along the left side of the heatmap indicates the sampling site for each colony.

#### Proteomic dataset

##### Montipora

Proteomic data was generated for 99 *Montipora* sp. samples, resulting in a raw host and symbiont dataset containing 10,606 *Montipora* proteins and 5,081 symbiont proteins. Only host-derived proteins were targeted for cleaning, however, symbiont proteins for all three coral genera are available in the raw datasets. After quality filtering to remove symbiont proteins and proteins with poor confidence (PEP and Q-value >0.01, Precursor Quantity > 1), 98 samples remained (one contained only symbiont proteins) and 9,510 host proteins. Before additional filtering, run intensities were generally consistent across samples and had a normal distribution (**Figs. 6A-B**). However, data completeness varied greatly by sample from 19.7% in M036 to 82.6% in M015 (**Table S10**) and samples with the lowest completeness clustered away from all other samples on PC1 (61.9% variation) and PC2 (10.9% variation; **Fig. 6C**). Protein missingness across the entire dataset with 46.71% and appeared lowest in MogMog Landing, Asor Landing and Falalop Men’s House and highest in Yealil Inside, Federai Landing and Sohng (**Fig. 6D**). After removing samples (M26.1, M34, M35, M36, M37, M100) and proteins with high missingness (>40%) in any site, 93 samples and 2,673 proteins remained (overall missingness 2.83%; **Fig. 6H**) for missing value imputation and downstream analyses. After normalization and log2-fold scaling, the log2 intensities all remaining proteins appeared normally distributed (**Fig. 6E-F)** and there were no visual outliers on PC1 (45.3% variation) and PC2 (5.4% variation) (**Fig. 6G**).

**Figure 6.**
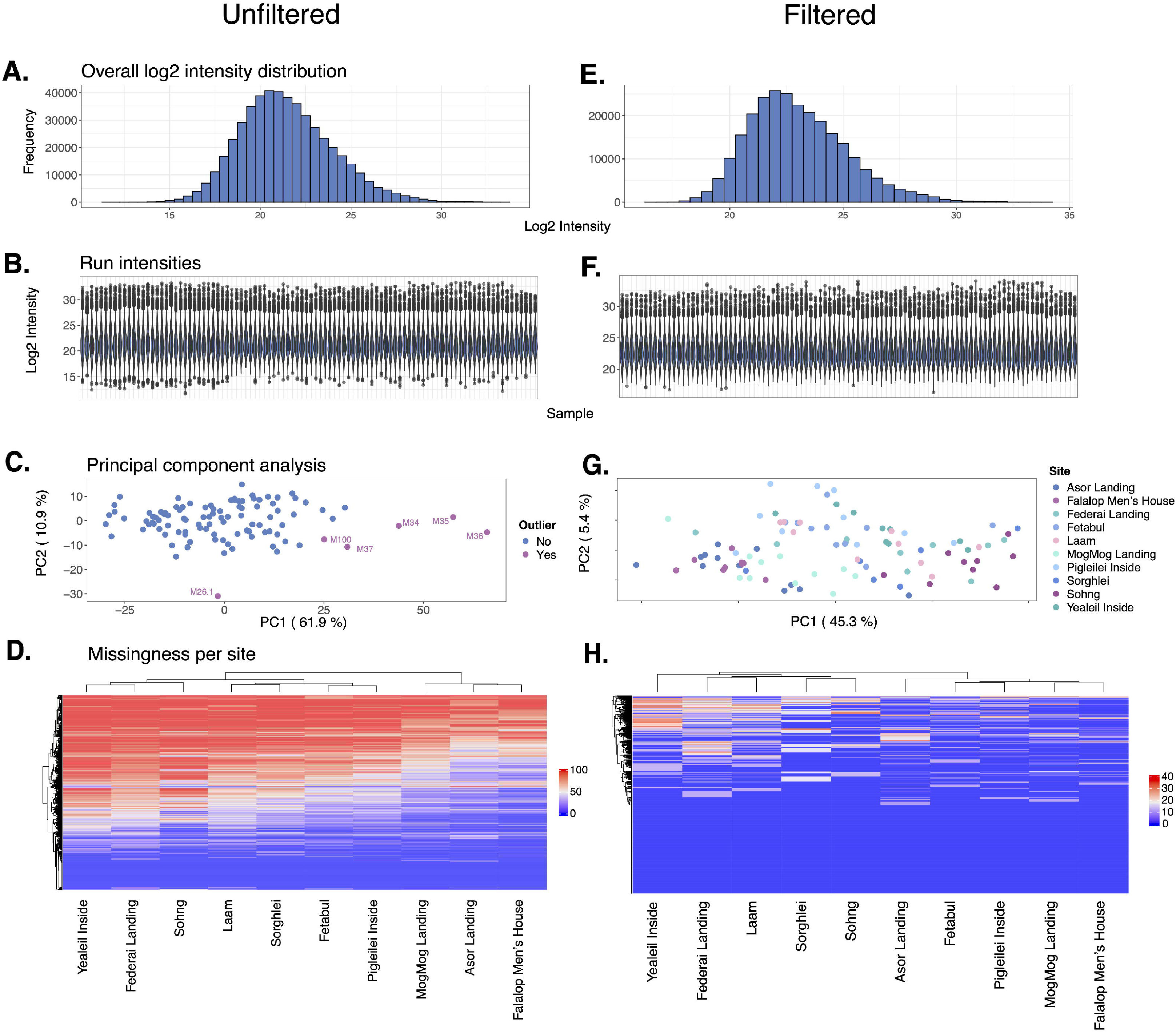
Quality control of *Montipora* proteomic data. (A-D) Metrics calculated before quality filtering and missing value imputation. **(E-H)** Metrics calculated after quality filtering and missing value imputation. (**A, E**) Distribution of log2 intensity for all protein groups across all samples. **(B, F)** Violin plot showing per-sample log2 intensity distributions. (**C, G**) Principal coordinates analysis based on normalized log2 intensity profiles of protein groups. In (**C**), colors correspond to outliers that were removed from the final dataset (violet) and those that were retained (periwinkle). In (**G**), colors correspond to collection site. (**D, H**) Heatmap of protein group missingness before missing value imputation across sites. Rows represent protein groups and columns represent collection sites. Cell color indicates the proportion of samples with missing data for each site. Samples are hierarchically clustered based on the missingness from each site.

##### Pocillopora

Proteomic data was generated for 25 *Pocillopora* spp. samples, resulting in a combined host and symbiont dataset containing 12,869 *Pocillopora* proteins and 7,860 symbiont proteins. Only host-derived proteins were retained for downstream cleaning. After quality filtering to remove symbiont proteins and proteins with poor confidence (PEP and Q-value >0.01, Precursor Quantity > 1), 11,679 host proteins remained. Before additional filtering, run intensities were largely consistent across samples and followed a normal distribution (**Fig. 7A-B**). Data completeness was consistent across all samples, ranging from 61.4% in P18 to 77.2% in P11 (**Table S10**). Samples collected from the same site tended to cluster together on PC1 (19.3% variation) and PC2 (12.8% variation) in the raw data PCA, except for samples P15 and P11, which grouped together with samples P207 and P208 from MogMog Landing (**Fig. 7C**). Overall protein missingness across the dataset was 28.67% and protein missingness was generally consistent across the three sites (**Fig. 7D**). After proteins with high missingness (>40%) across all sites, 6,594 proteins remained (2.52% overall missingness) for missing value imputation and downstream analyses **(Fig. 7H**). Following normalization and log2-fold scaling, the log2 intensities of all remaining proteins were normally distributed (**Figs. 7E-F**), and there were no clear outliers on PC1 (15.5% variation) and PC2 (13.3% variation) (**Fig. 7G**).

**Figure 7.**
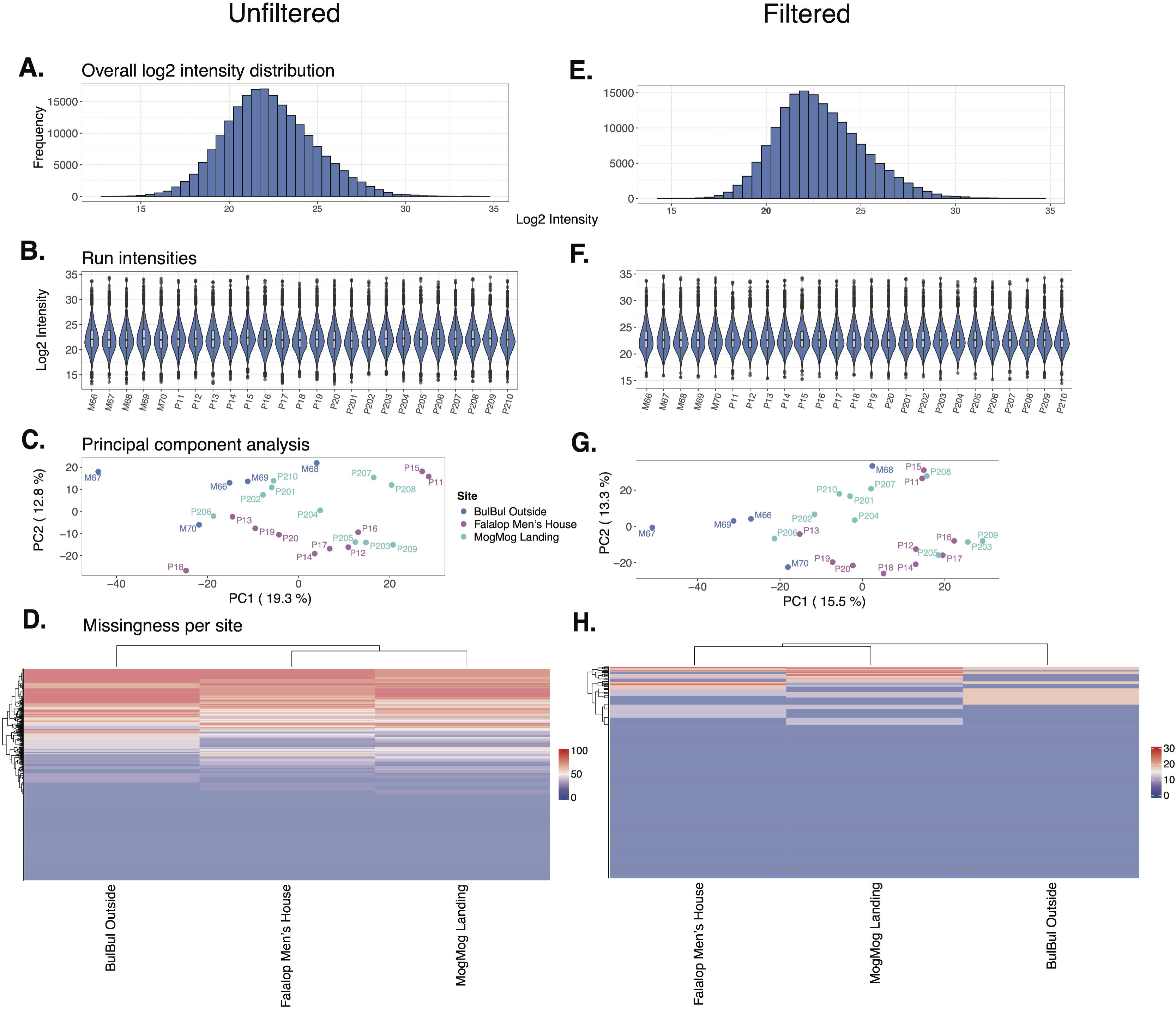
Quality control of *Pocillopora* proteomic data. (A-D) Metrics calculated before quality filtering and missing value imputation. **(E-H)** Metrics calculated after quality filtering and missing value imputation. (**A, E**) Distribution of log2 intensity for all protein groups across all samples. **(B, F)** Violin plot showing per-sample log2 intensity distributions. (**C, G**) Principal coordinates analysis based on normalized log2 intensity profiles of protein groups. Colors correspond to collection site. (**D, H**) Heatmap of protein group missingness across sites. Rows represent protein groups and columns represent collection sites. Cell color indicates the proportion of samples with missing data before missing value imputation for each site. Samples are hierarchically clustered based on the missingness from each site.

##### Acropora

Proteomic data was generated for 25 *Acropora* spp. samples, resulting in a combined host and symbiont dataset containing 13,081 *Acropora* proteins and 6,845 symbiont proteins. Although symbiont proteins are included in the raw dataset, only host proteins were included in the final, cleaned dataset. After quality filtering to remove symbiont proteins and proteins with poor confidence (PEP and Q-value >0.01, Precursor Quantity > 1), 11,613 host proteins remained. The run intensities of the raw dataset followed a normal distribution, and the run intensities were consistent across samples (**Figs. 8A-B**). Data completeness ranged from 53.6% in A338 to 74.2% in A27 and A309 (**Table S10**). Samples collected from the MogMog Landing clustered separately from samples collected from BulBul Outside and Falalop Men’s House on PC2 (11.1% variation) and there were no visual outliers on either PC1 (27.9% variation) or PC2 **(Fig. 8A**). Overall protein missingness across the dataset was 30.02% and protein missingness was consistent across the three sites (**Fig. 8D**). After proteins with high missingness (>40%) across all sites, 6,254 proteins remained (3.20% overall missingness) for missing value imputation and downstream analyses **(Fig. 8H**). Following normalization and log2-fold scaling, the log2 intensities of all remaining proteins were normally distributed (**Figs. 8E-F**), and there were no clear outliers on PC1 (21.3% variation) and PC2 (13.1% variation) (**Fig. 8H**).

**Figure 8.**
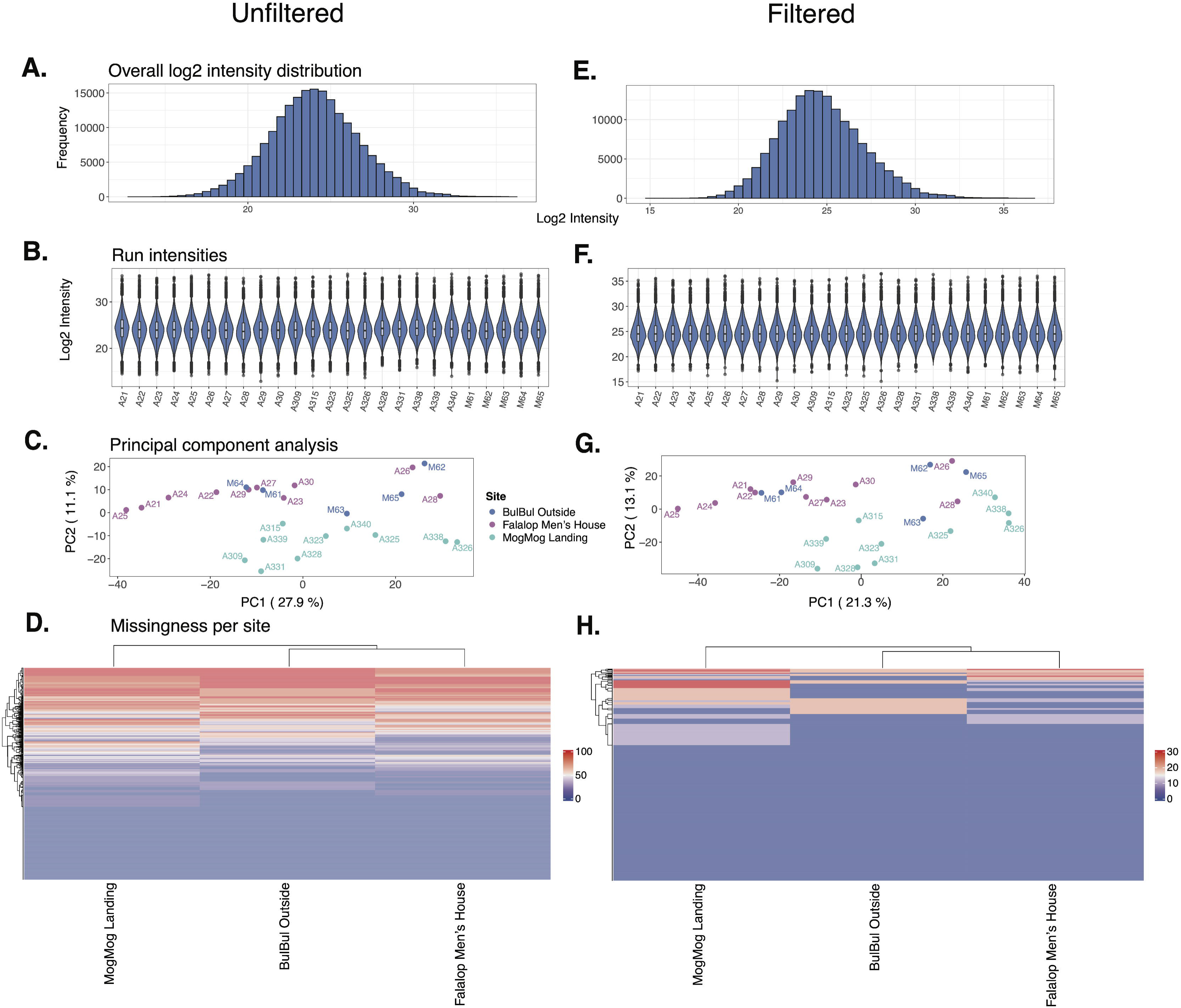
Quality control of *Acropora* proteomic data. (A-D) Metrics calculated before quality filtering and missing value imputation. **(E-H)** Metrics calculated after quality filtering and missing value imputation. (**A, E**) Distribution of log2 intensity for all protein groups across all samples. **(B, F)** Violin plot showing per-sample log2 intensity distributions. (**C, G**) Principal coordinates analysis based on normalized log2 intensity profiles of protein groups. Colors correspond to collection site. (**D, H**) Heatmap of protein group missingness across sites. Rows represent protein groups and columns represent collection sites. Cell color indicates the proportion of samples with missing data before missing value imputation for each site. Samples are hierarchically clustered based on the missingness from each site.

**Figure 9.**
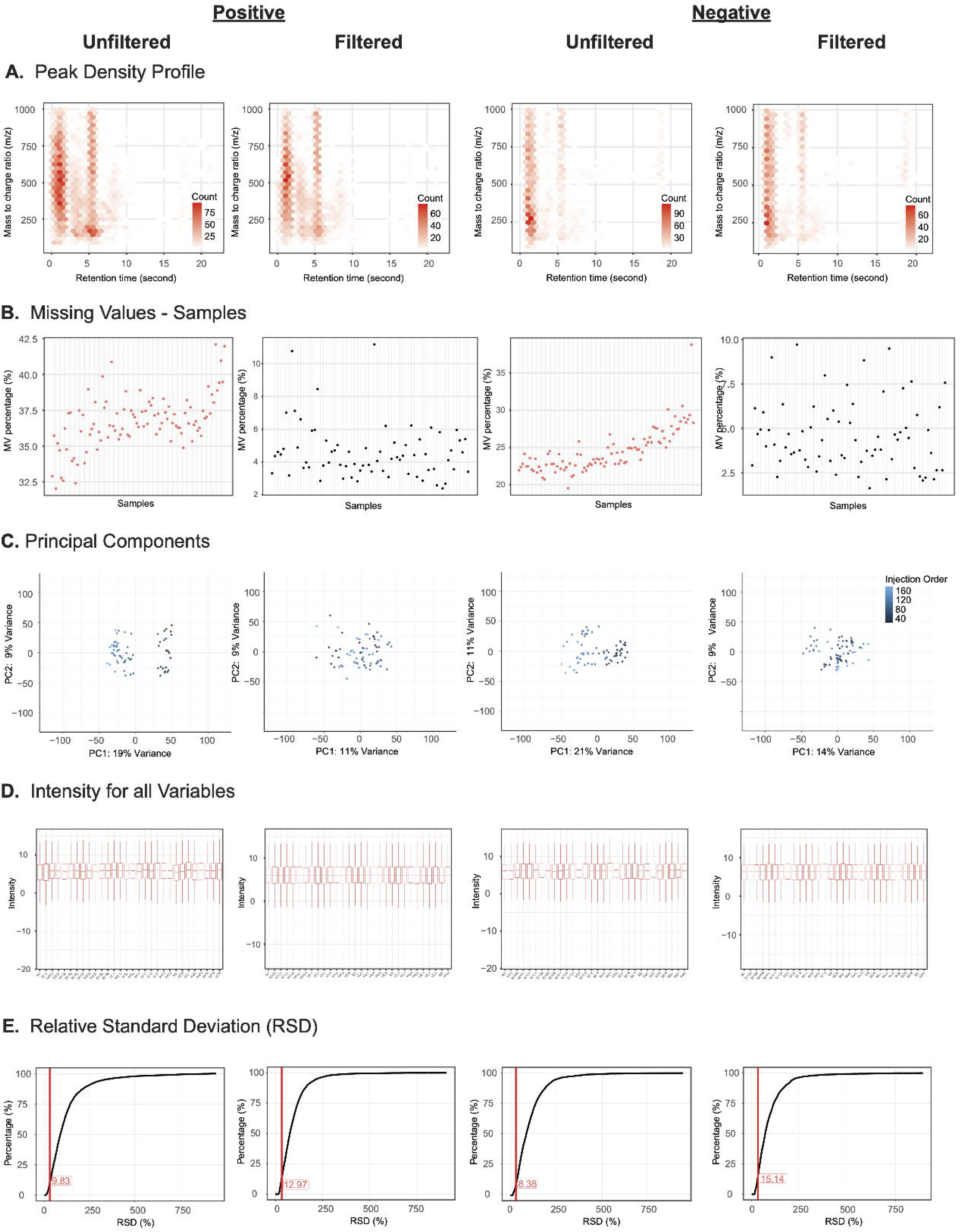
Quality control of *Montipora* metabolomic data. **(A)** Density of metabolites identified across the spectrum of mass-to-charge ratio and retention time, across (from left to right) positive ionization unfiltered and filtered features and negative ionization unfiltered and filtered features. **(B)** MV percentage represents the percentage of metabolites detected in a given sample relative to the total set of metabolites identified for all samples within that genus. Samples are ordered by injection order (**Table S4**). **(C)** Principal coordinates analysis based on log2 intensity profiles of metabolites. Colors correspond to injection order. **(D)** Per-sample distribution of metabolite log2 intensities. Positive samples are in batch 3, negative samples are in batch 4. Only first 30 samples shown. **(E)** The cumulative distribution of relative standard deviation (RSD) across detected metabolites. The red line indicates 30% RSD.

#### Metabolomic dataset

##### Montipora

LC-MS data was generated for 97 *Montipora* spp. samples, resulting in 8,623 positive ionization mode metabolites (36.72% missingness overall) and 4,310 negative ionization mode metabolites (24.49% missingness overall; **Fig. 9A**, **Fig. 9D**). Metabolites were filtered for missingness, leaving 5,205 positive ionization mode metabolites and 3,183 negative ionization mode metabolites (**Fig. 9B**). We detected 12 outliers in the positive ionization mode data and 15 outliers in the negative ionization mode data, which we removed from both positive and negative datasets (**Fig. 9B**), leaving 70 samples in both positive (4.65% missingness overall) and negative (4.58% missingness overall) ionization modes for missing value imputation. PCA sample clustering (**Fig. 9C)** indicated that injection order had a large effect on the global variance and structure, so we removed this effect using *removeBatchEffect* (**Fig. 9C**). We did not identify any additional samples as outliers on the PCA (**Fig. 9C**). Relative standard deviation (RSD) changed in both the positive (9.83 to 12.97) and negative ionization modes (8.38 to 15.14), yet the structure of the plots did not change, indicating successful removal of outliers and technical noise (**Fig. 9E**).

##### Pocillopora

LC-MS data was generated for 24 *Pocillopora* spp. samples, resulting in 8,623 positive ionization mode metabolites (38.94% missingness overall) and 4,310 negative ionization mode metabolites (26.99% missingness overall) (**Fig. 10A**, **Fig. 10D**). Metabolites were filtered for missingness, leaving 4,927 positive ionization mode metabolites and 3,100 negative ionization mode metabolites (**Fig. 10B**). We did not detect any outliers; random forest imputation was run on all 24 samples for both positive (7.84% missingness overall) and negative (4.74% missingness overall) ionization modes. PCA sample clustering and per-sample missingness indicated that global variance was strongly associated with injection order, so we removed this effect using *removeBatchEffect* (**Fig. 10C**). We identified sample P14 as an outlier based on negative ionization mode PCA clustering and removed it from both datasets **(Fig. 10C)**. Relative standard deviation (RSD) changed for both the positive (4.63 to 2.84) and negative ionization modes (8.89 to 10.81). However, the structure of the plots was unchanged, indicating successful removal of outliers and technical noise (**Fig. 10E**).

**Figure 10.**
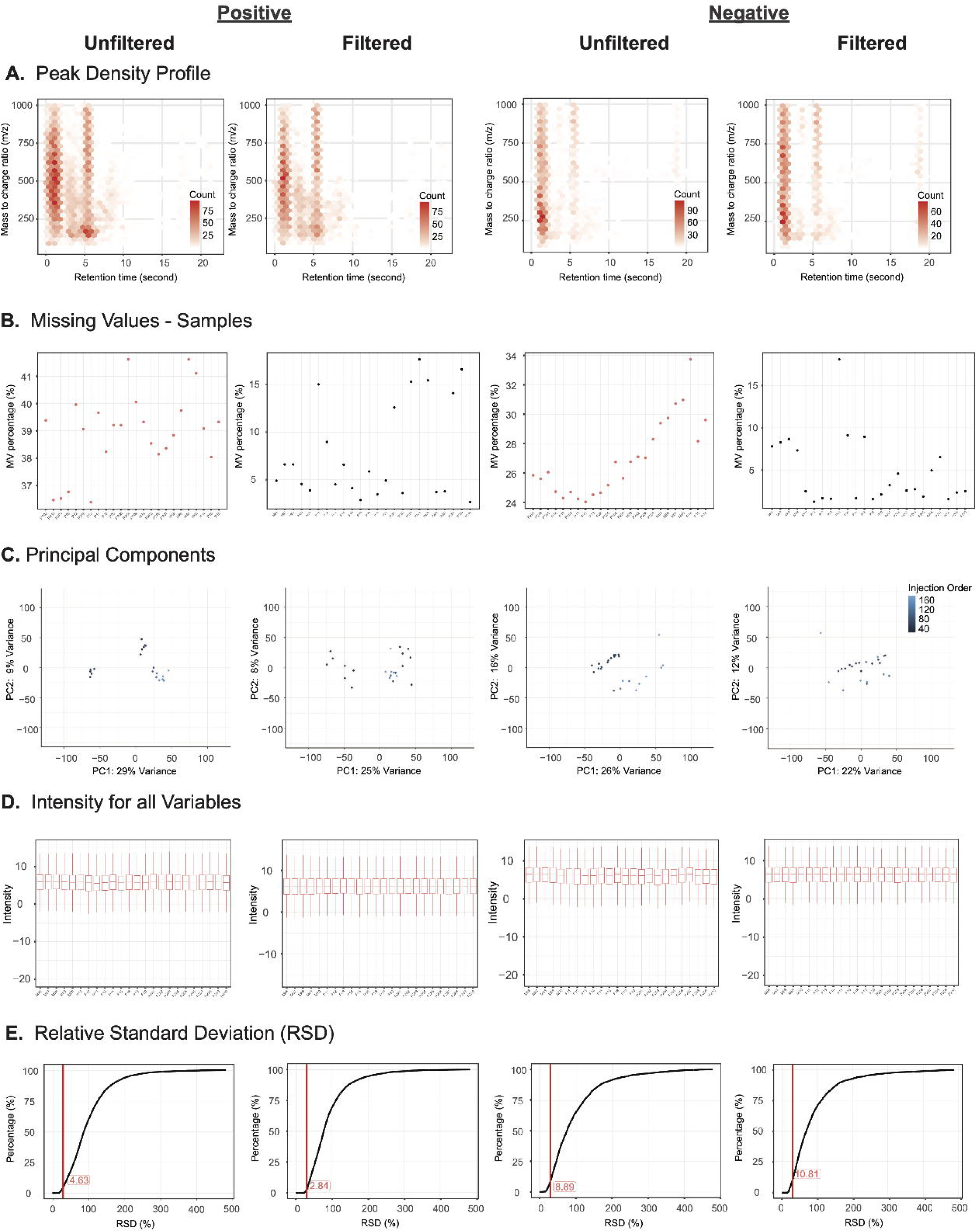
Quality control of *Pocillopora* metabolomic data. **(A)** Density of metabolites identified across the spectrum of mass-to-charge ratio and retention time, across (from left to right) positive ionization unfiltered and filtered features and negative ionization unfiltered and filtered features. **(B)** MV percentage represents the percentage of metabolites detected in a given sample relative to the total set of metabolites identified for all samples within that genus. Samples are ordered by injection order. **(C)** Principal coordinates analysis based on log2 intensity profiles of metabolites. Colors correspond to injection order. **(D)** Per-sample distribution of metabolite log2 intensities. Positive samples are in batch 3, negative samples are in batch 4. Only first 30 samples shown. **(E)** The cumulative distribution of relative standard deviation (RSD) across detected metabolites. The red line indicates 30% RSD.

**Figure 11.**
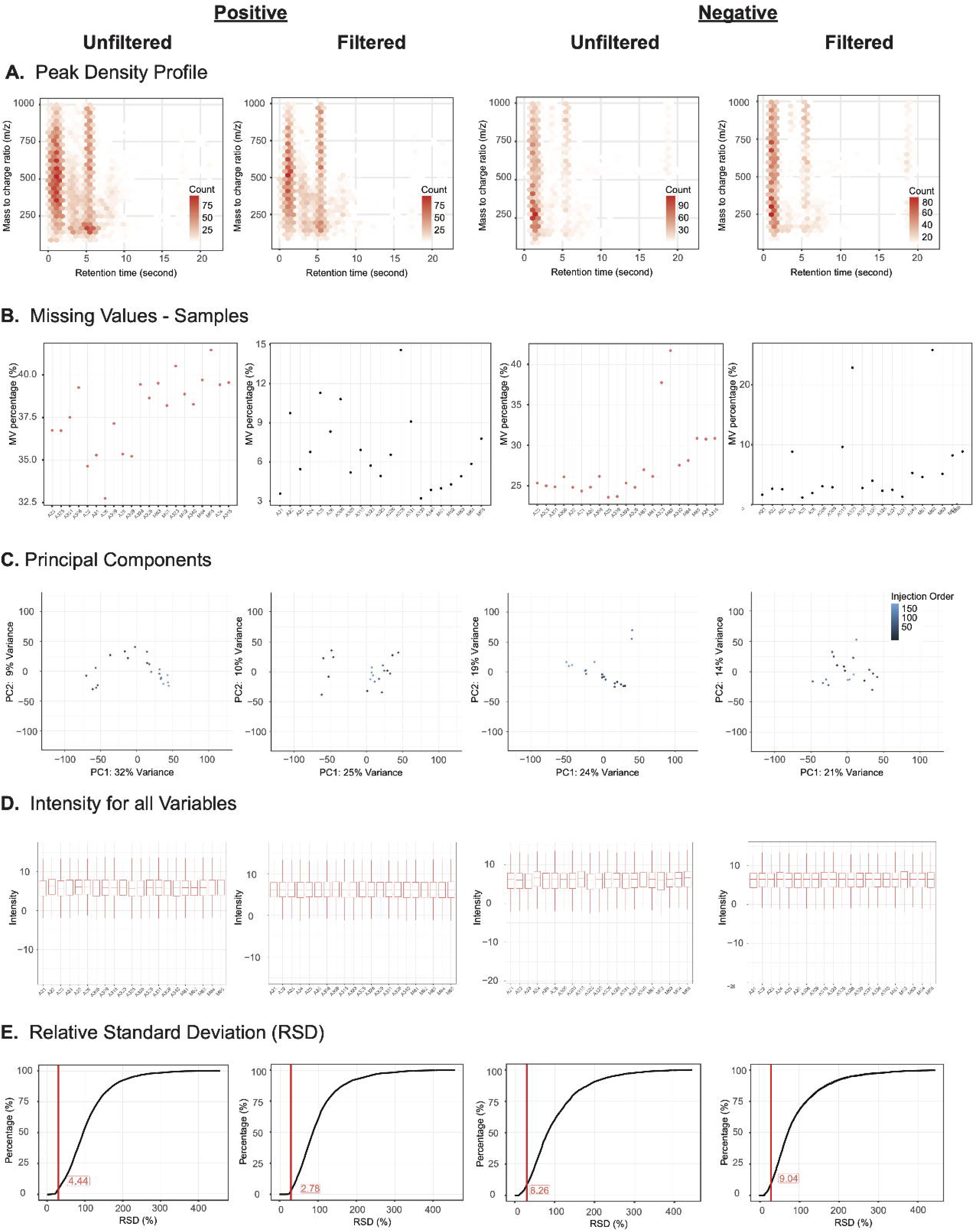
Quality control of *Acropora* metabolomic data. **(A)** Density of metabolites identified across the spectrum of mass-to-charge ratio and retention time, across (from left to right) positive ionization unfiltered and filtered features and negative ionization unfiltered and filtered features. **(B)** MV percentage represents the percentage of metabolites detected in a given sample relative to the total set of metabolites identified for all samples within that genus. Samples are ordered by injection order. **(C)** Principal coordinates analysis based on log2 intensity profiles of metabolites. Colors correspond to injection order. **(D)** Per-sample distribution of metabolite log2 intensities. Positive samples are in batch 3, negative samples are in batch 4. Only first 30 samples shown. **(E)** The cumulative distribution of relative standard deviation (RSD) across detected metabolites. The red line indicates 30% RSD.

##### Acropora

LC-MS data was generated for 21 *Acropora* spp. samples, resulting in 8,623 positive (37.82% missingness overall) and 4,310 negative (27.60% missingness overall; **Fig. 11A**, **Fig. 11D**) ionization mode metabolites. Metabolites were filtered for missingness leaving 4,959 positive and 3,097 negative ionization mode metabolites (**Fig. 11B**). All 21 samples were used for missing value imputation for both positive (6.79% missingness overall) and negative (6.11% missingness overall) ionization modes. Per-sample missingness and PCA sample clustering indicated that injection order has an outsized effect on the overall variance and structure, so we removed this effect using *removeBatchEffect* (**Fig. 11C**). We identified samples M62 and A323 as outliers based on negative ionization mode PCA clustering and removed it from both datasets **(Fig. 11C)**. Relative standard deviation (RSD) changed for both positive (4.44 to 2.78) and negative (8.26 to 9.04) ionization modes, but the structure of the plots did not change, indicating successful removal of outliers and technical noise (**Fig. 11E**).

### Re-use potential

The datasets described here provide a multi-omics resource for investigating how environmental conditions and human activity influence coral molecular phenotypes. By integrating genomic, proteomic, and metabolomic data from coral colonies sampled across an environmental gradient, these data can be used to investigate anthropogenic disturbance effects on the coral holobiont across different omics layers. Because reefs at Ulithi have been previously classified into ecological clusters associated with differing degrees of human presence and environmental exposure [8] (**Figure 1**, **Table 1**), this dataset provides an opportunity to investigate how omics data reflect coral phenotypes and how they vary across well-characterized gradients of human impact.

A major goal in coral conservation is to build a suite of low-cost, point-of-care monitoring tools that are diagnostic or predictive of coral health to facilitate coral conservation efforts. Some of these innovative tools rely on the generation of integrated multi-omics information, in particular transcriptomics, proteomics, and metabolomics data, to identify and validate biomarkers that can be readily integrated into existing point-of-care testing platforms, such as lateral flow tests and colorimetric dipstick tests [12]. To date, most putative coral stress biomarkers are diagnostic of thermal stress [12,39]. Additional work is needed to identify and validate biomarkers that are either predictive of resilience or adaptive capacity or are diagnostic of the broad range of stressors faced by corals. As stress is a major threat to coral health, there is significant interest in biomarkers to elucidate chronic stress from anthropogenic activities.

A key challenge in biomarker discovery and validation is understanding what drives variation in multi-omics datasets. Corals exist in diverse environmental settings, harbor different symbionts, and exhibit substantial intraspecific genetic variation. Therefore, it is imperative to discover biomarkers that show consistent behavior across genotypic, environmental, and holobiont variability [12]. Although much progress has been made in accounting for these effects in gene expression data, suitable biomarkers are yet to be identified that address proteome and metabolome variation [12,40]. These data can be used as a resource to identify biomarkers associated with local human activities to assess their utility (e.g., consistency) in different reefs and environmental conditions. Because samples include multiple coral genera (*Montipora*, *Pocillopora*, and *Acropora*), our datasets also enable comparative analysis among coral taxa.

As coral reefs increasingly experience combined pressures from climate change and local anthropogenic stressors [5–7], datasets that integrate ecological context with genomic, proteomic, and metabolomic data will be critical for understanding the molecular underpinnings of coral resilience. The resources presented here therefore provide a platform for future work linking coral molecular biology with ecosystem-level change and for developing molecular biomarkers that can facilitate evidence-based ecosystem management practices.

## Data Availability

Raw and clean high coverage whole-genome variant (*Montipora* only), proteomic, and metabolomic datasets for *Montipora*, *Pocillopora*, and *Acropora* colonies collected from Ulithi Atoll, Yap State, FSM in June 2023, along with photographs of sampling sites and sampled colonies are available on Zenodo (https://zenodo.org/records/19582294). Raw sequencing reads for *Montipora* are available from NCBI’s SRA repository (BioProject <u>PRJNA1121902</u>). Metadata, scripts, and output related to dataset generation and QC are available from https://github.com/echille/Ulithi_2023_Coral_Multi-Omic_Dataset.

## Abbreviations

WGS: Whole Genome Sequencing
MAF: Minor Allele Frequency
RIPA: Radio-Immunoprecipitation Assay
LC-MS: Liquid Chromatography–Mass Spectrometry
DIA: Data Independent Acquisition
PEP: Posterior Error Probability
PCA: Principal Component Analysis
SD: Standard Deviation
SNP: Single Nucleotide Polymorphisms
RSD: Relative Standard Deviation

## Author’s Contributions

Conceptualization: DB NLC MP JR GB. Data curation: EEC GMP.

Formal analysis: EEC GMP TGS. Funding acquisition: DB NLC MP JR GB. Investigation: NLC MP PN AA GB. Methodology: EEC.

Supervision: DB NLC MP JR. Visualization: EEC GMP.

Writing – original draft: EEC GMP.

Writing – review & editing: EEC GMP TGS DB NLC MP JR GB.

## Acknowledgments

We gratefully acknowledge the people of Ulithi, Federated States of Micronesia, for permission and their assistance in collecting coral samples, and for their continued collaboration in the studies of their Atoll. We also thank Dr. Haiyan Zheng of the Biological Mass Spectrometry Facility of Robert Wood Johnson Medical School and Rutgers, The State University of New Jersey for generating the proteomic dataset as well as Dr. Sonam Tamrakar, Dr. Jeremy L. Balsbaugh, and Dr. Jennifer C. Liddle of the UConn Proteomics & Metabolomics Facility for conducting the untargeted metabolomics analysis.

## References

1. Mcleod E, Anthony KRN, Mumby PJ, Maynard J, Beeden R, Graham NAJ, et al.. The future of resilience-based management in coral reef ecosystems. J Environ Manage. 2019; doi: 10.1016/j.jenvman.2018.11.034.

2. Voolstra C, Miller D, Ragan M, Hoffmann A, Hoegh-Guldberg O, Bourne D, et al.. The ReFuGe 2020 Consortium—using “omics” approaches to explore the adaptability and resilience of coral holobionts to environmental change. Frontiers in Marine Science. 2015; doi: 10.3389/fmars.2015.00068.

3. Woodhead AJ, Hicks CC, Norström AV, Williams GJ, Graham NAJ. Coral reef ecosystem services in the Anthropocene. Funct Ecol. Wiley; 2019; doi: 10.1111/1365-2435.13331.

4. Schlichter D, Zscharnack B, Krisch H. Transfer of photoassimilates from endolithic algae to coral tissue. Sci Nat. Springer Science and Business Media LLC; 1995; doi: 10.1007/s001140050234.

5. Carpenter KE, Abrar M, Aeby G, Aronson RB, Banks S, Bruckner A, et al.. One-third of reef-building corals face elevated extinction risk from climate change and local impacts. Science. 2008; doi: 10.1126/science.1159196.

6. Donovan MK, Burkepile DE, Kratochwill C, Shlesinger T, Sully S, Oliver TA, et al.. Local conditions magnify coral loss after marine heatwaves. Science.

7. National Academies of Sciences, Engineering, and Medicine. A Research Review of Interventions to Increase the Persistence and Resilience of Coral Reefs. Washington, D.C., DC: The National Academies Press;

8. Crane NL, Nelson P, Abelson A, Precoda K, Rulmal J Jr, Bernardi G, et al.. Atoll-scale patterns in coral reef community structure: Human signatures on Ulithi Atoll, Micronesia. PLoS One. 2017; doi: 10.1371/journal.pone.0177083.

9. Cannon SE, Aram E, Beiateuea T, Kiareti A, Peter M, Donner SD. Coral reefs in the Gilbert Islands of Kiribati: Resistance, resilience, and recovery after more than a decade of multiple stressors. PLoS One. 2021; doi: 10.1371/journal.pone.0255304.

10. Brown KT, Bender-Champ D, Bryant DEP, Dove S, Hoegh-Guldberg O. Human activities influence benthic community structure and the composition of the coral-algal interactions in the central Maldives. J Exp Mar Bio Ecol. Elsevier BV; 2017; doi: 10.1016/j.jembe.2017.09.006.

11. Muthukrishnan R, Fong P. Multiple anthropogenic stressors exert complex, interactive effects on a coral reef community. Coral Reefs. Springer Science and Business Media LLC; 2014; doi: 10.1007/s00338-014-1199-1.

12. Chille EE, Stephens TG, Nandi S, Jiang H, Gerdes MJ, Williamson OM, et al.. Coral restoration in the omics era: Development of point-of-care tools for monitoring disease, reproduction, and thermal stress. Bioessays. Wiley; 2025; doi: 10.1002/bies.70007.

13. Bernardi G, Gatins R, Paddack M, Nelson P, Rulmal J, Crane N. Genomics of a novel ecological phase shift: the case of a ‘weedy’ Montipora coral in Ulithi, Micronesia. Coral Reefs. 2024; doi: 10.1007/s00338-024-02486-9.

14. Crane N, Paddack M, Nelson P, Abelson A, Rulmal J, Bernardi G. Corallimorph and Montipora Reefs in Ulithi Atoll, Micronesia: documenting unusual reefs. J Ocean Sci Found. 2016; doi: 10.5281/zenodo.51289.

15. Stephens TG, Lee J, Jeong Y, Yoon HS, Putnam HM, Majerová E, et al.. High-quality genome assembles from key Hawaiian coral species. Gigascience. 2022; doi: 10.1093/gigascience/giac098.

16. Poplin R, Chang P-C, Alexander D, Schwartz S, Colthurst T, Ku A, et al.. A universal SNP and small-indel variant caller using deep neural networks. Nat Biotechnol. Springer Science and Business Media LLC; 2018; doi: 10.1038/nbt.4235.

17. Chen S. Ultrafast one pass FASTQ data preprocessing, quality control, and deduplication using fastp. Imeta. Wiley; 2023; doi: 10.1002/imt2.107.

18. Andrews S. FastQC: a quality control tool for high throughput sequence data. Babraham Bioinformatics, Babraham Institute, Cambridge, United Kingdom;

19. Ewels P, Magnusson M, Lundin S, Käller M. MultiQC: summarize analysis results for multiple tools and samples in a single report. Bioinformatics. 2016; doi: 10.1093/bioinformatics/btw354.

20. Vasimuddin M, Misra S, Li H, Aluru S. Efficient architecture-aware acceleration of BWA-MEM for multicore systems. 2019 IEEE International Parallel and Distributed Processing Symposium (IPDPS). IEEE;

21. Danecek P, Bonfield JK, Liddle J, Marshall J, Ohan V, Pollard MO, et al.. Twelve years of SAMtools and BCFtools. Gigascience. Oxford University Press (OUP); 2021; doi: 10.1093/gigascience/giab008.

22. Okonechnikov K, Conesa A, García-Alcalde F. Qualimap 2: advanced multi-sample quality control for high-throughput sequencing data. Bioinformatics. Oxford University Press (OUP); 2016; doi: 10.1093/bioinformatics/btv566.

23. Korunes KL, Samuk K. pixy: Unbiased estimation of nucleotide diversity and divergence in the presence of missing data. Mol Ecol Resour. Wiley; 2021; doi: 10.1111/1755-0998.13326.

24. Genomics in the Cloud: Analyzing Next-Generation Sequencing Data with Docker and GATK.

25. Zhang J, Richards ZT, Adam AAS, Chan CX, Shinzato C, Gilmour J, et al.. Evolutionary responses of a reef-building coral to climate change at the end of the last glacial maximum. Mol Biol Evol. Oxford University Press (OUP); 2022; doi: 10.1093/molbev/msac201.

26. Hughes CS, Moggridge S, Müller T, Sorensen PH, Morin GB, Krijgsveld J. Single-pot, solid-phase-enhanced sample preparation for proteomics experiments. Nat Protoc. Springer Science and Business Media LLC; 2019; doi: 10.1038/s41596-018-0082-x.

27. Demichev V, Messner CB, Vernardis SI, Lilley KS, Ralser M. DIA-NN: neural networks and interference correction enable deep proteome coverage in high throughput. Nat Methods. Springer Science and Business Media LLC; 2020; doi: 10.1038/s41592-019-0638-x.

28. Fuller ZL, Mocellin VJL, Morris LA, Cantin N, Shepherd J, Sarre L, et al.. Population genetics of the coral Acropora millepora: Toward genomic prediction of bleaching. Science. American Association for the Advancement of Science (AAAS); 2020; doi: 10.1126/science.aba4674.

29. Buitrago-López C, Mariappan KG, Cárdenas A, Gegner HM, Voolstra CR. The Genome of the Cauliflower Coral Pocillopora verrucosa. Genome Biol Evol. Oxford University Press (OUP); 2020; doi: 10.1093/gbe/evaa184.

30. Chen Y, Shah S, Dougan KE, van Oppen MJH, Bhattacharya D, Chan CX. Improved Cladocopium goreaui genome assembly reveals features of a facultative coral symbiont and the complex evolutionary history of dinoflagellate genes. Microorganisms. MDPI AG; 2022; doi: 10.3390/microorganisms10081662.

31. Dougan KE, Bellantuono AJ, Kahlke T, Abbriano RM, Chen Y, Shah S, et al.. Whole-genome duplication in an algal symbiont bolsters coral heat tolerance. Sci Adv. American Association for the Advancement of Science (AAAS); 2024; doi: 10.1126/sciadv.adn2218.

32. Quast J-P, Schuster D, Picotti P. protti: an R package for comprehensive data analysis of peptide- and protein-centric bottom-up proteomics data. Bioinform Adv. Oxford University Press (OUP); 2022; doi: 10.1093/bioadv/vbab041.

33. Stekhoven DJ, Bühlmann P. MissForest--non-parametric missing value imputation for mixed-type data. Bioinformatics. Oxford University Press (OUP); 2012; doi: 10.1093/bioinformatics/btr597.

34. Smith CA, Maille GO, Want EJ, Qin C, Trauger SA, Brandon TR, et al.. METLIN. Ther Drug Monit. Ovid Technologies (Wolters Kluwer Health); 2005; doi: 10.1097/01.ftd.0000179845.53213.39.

35. Shen X, Yan H, Wang C, Gao P, Johnson CH, Snyder MP. TidyMass an object-oriented reproducible analysis framework for LC-MS data. Nat Commun. Springer Science and Business Media LLC; 2022; doi: 10.1038/s41467-022-32155-w.

36. Smilde AK, van der Werf MJ, Bijlsma S, van der Werff-van der Vat BJC, Jellema RH. Fusion of mass spectrometry-based metabolomics data. Anal Chem. American Chemical Society (ACS); 2005; doi: 10.1021/ac051080y.

37. Ritchie ME, Phipson B, Wu D, Hu Y, Law CW, Shi W, et al.. Limma powers differential expression analyses for RNA-sequencing and microarray studies. Nucleic Acids Res. Oxford University Press; 2015; doi: 10.1093/nar/gkv007.

38. Parkinson JE, Baker AC, Baums IB, Davies SW, Grottoli AG, Kitchen SA, et al.. Molecular tools for coral reef restoration: beyond biomarker discovery. Conservation Letters. Wiley Online Library; 2020; doi: 10.1111/conl.12687.

39. Bhattacharya D, Nandi S, Chille EE, Arroyo M, Stephens TG. The host Coral Bleaching response viewed through the lens of multi-omics: Multi-omics provides the tools to understand the complex molecular basis of Coral Bleaching, which can aid conservation efforts. Bioessays. Wiley; 2026; doi: 10.1002/bies.70110.

